# Structural and molecular basis for Cardiovirus 2A protein as a viral gene expression switch

**DOI:** 10.1101/2020.08.11.245035

**Authors:** Chris H. Hill, Sawsan Napthine, Lukas Pekarek, Anuja Kibe, Andrew E. Firth, Stephen C. Graham, Neva Caliskan, Ian Brierley

**Author notes:** authors contributed equally to this work (Chris H. Hill, Sawsan Napthine, Lukas Pekarek).

## Abstract

Programmed −1 ribosomal frameshifting (PRF) in cardioviruses is activated by the 2A protein: a multi-functional virulence factor that also inhibits cap-dependent translational initiation. Here we present the X-ray crystal structure of 2A and show that it selectively binds to and stabilises the PRF stimulatory RNA element in the viral genome. Using optical tweezers, we define the conformational repertoire of this element and measure changes in unfolding pathways arising from mutation and 2A binding. Next, we demonstrate a strong interaction between 2A and the small ribosomal subunit and present a cryo-EM structure of 2A bound to initiated 70S ribosomes. Multiple copies of 2A bind to the 16S rRNA where they may compete for binding with initiation and elongation factors. Together, these results define the structural basis for RNA recognition by 2A, expand our understanding of the 2A-dependent gene expression switch and reveal how 2A accumulation may shut down translation during virus infection.

## Introduction

Encephalomyocarditis virus (EMCV) is the archetype of the *Cardiovirus A* group within the family *Picornaviridae*. It has a 7.8 kb positive-sense, single-stranded, linear RNA genome comprising a single long open reading frame (ORF; ~2200 amino acids) flanked by an extended 5′ untranslated region (UTR) containing an internal ribosome entry site (IRES), and a shorter 3′ UTR with a poly(A) tail. Upon infection, the genome is translated directly to yield a polyprotein (L-1ABCD-2ABC-3ABCD) that is proteolytically processed into approximately 12 individual gene products by the viral 3C protease. In addition to IRES utilisation (Jang et al., 1988), the discovery of Stop-Go peptide release (Hahn and Palmenberg, 2001; Palmenberg et al., 1992) and PRF (Loughran et al., 2011; Napthine et al., 2017) during genome translation has established EMCV as a model system for studying ribosome-related gene expression mechanisms. In EMCV, PRF occurs 11–12 codons into the start of the 2B gene, with up to 70% of ribosomes changing frame and producing the 2B* *trans*-frame product.

PRF is a translational control strategy employed by many RNA viruses, where it ensures the production of proteins in optimal ratios for efficient virus assembly, and enables viruses to expand their coding capacity through the utilisation of overlapping ORFs (reviewed in Atkins et al., 2016; Firth and Brierley, 2012; Korniy et al., 2019). In canonical PRF, elongating ribosomes pause over a heptanucleotide “slippery sequence” of the form X_XXY_YYZ when they encounter a “stimulatory element” 5–9 nucleotides downstream in the mRNA. During this time, a –1 frameshift may occur if codon-anticodon re-pairing takes place over the X_XXY_YYZ sequence: the homopolymeric stretches allow the tRNA in the P-site tRNA to slip from XXY to XXX, and the tRNA in the A-site to slip from YYZ to YYY. Frameshifting may occur during a late stage of the EF-G/eEF2 catalysed translocation step, with the stimulatory element causing paused ribosomes to become trapped in a chimeric rotated or hyper-rotated state that is relieved by either the spontaneous unfolding of the blockade or a –1 slip on the mRNA (Caliskan et al., 2014; Chen et al., 2014; Choi et al., 2020; Namy et al., 2006). A diverse array of stem-loops and pseudoknots are known to induce frameshifting, and the stability and unfolding kinetics of these stimulatory elements are thought to be the primary determinants of PRF efficiency (Chen et al., 2009; Giedroc and Cornish, 2009). More recently, the conformational plasticity of the elongation blockade has also been revealed to play an important role (Halma et al., 2019; Ritchie et al., 2012; Ritchie et al., 2014). Cardioviruses present a highly unusual variation to conventional viral PRF: the virally-encoded 2A protein is required as an essential *trans*-activator (Loughran et al., 2011), and the stimulatory element is thought to comprise an RNA-protein complex formed between 2A and a stem-loop in the viral RNA (Napthine et al., 2017). This unique mechanism allows for temporal control of gene expression as the efficiency of –1 frameshifting is linked to 2A concentration, which increases with time throughout the infection cycle (Napthine et al., 2017).

EMCV 2A is a small, basic protein (~17 kDa; 143 amino acids; pI ~9.1) generated by 3C-mediated proteolytic cleavage at the N-terminus (Jackson, 1986) and Stop-Go peptide release at a C-terminal 18-amino acid consensus sequence (Hahn and Palmenberg, 2001). Despite the identical name, the cardiovirus 2A has no homology to any other picornavirus “2A” protein (Yang et al., 2017), nor any other protein of known structure. Surprisingly, although cardiovirus replication and assembly is entirely cytoplasmic, 2A localises to nucleoli from early time-points post-infection (Aminev et al., 2003a, b). As well as its role in stimulating PRF (Napthine et al., 2017), 2A binds to 40S ribosomal subunits (Groppo and Palmenberg, 2007), inhibits apoptosis (Carocci et al., 2011) and contributes to host cell shut-off by inhibiting cap-dependent translation, despite having no protease activity against eIFs (Mosenkis et al., 1985). A previous mutational analysis (Groppo et al., 2011) identified a putative nuclear localisation sequence (NLS) of the form [G/P](K/R_3_)x_1-4_[G/P], similar to those found in yeast ribosomal proteins. This study also identified a C-terminal YxxxxLΦ motif, proposed to bind to and sequester eIF4E in a manner analogous to eIF4E binding protein 1 (4E-BP1), thereby inhibiting initiation of cap-dependent translation by interfering with eIF4F assembly (Merrick, 2015). Despite these insights, the absence of structural data has precluded a more definitive molecular characterisation of this multifunctional protein, and the mechanism by which it recognises RNA elements remains obscure.

Our previous RNA structure probing experiments suggested that the stimulatory element in the EMCV genome adopts a stem-loop conformation, and we have demonstrated that a conserved CCC motif in the putative loop region is essential for both 2A binding and PRF (Napthine et al., 2017). However, the nature of these interactions and the conformational dynamics of this frameshifting RNA element, including changes associated with 2A binding, are not well understood. To better define the conformational landscape of such RNAs, observing the folding and unfolding trajectories of individual molecules under tension can provide information beyond the resolution of conventional ensemble techniques, which are necessarily limited by molecular averaging (Visscher, 2016). In recent years, the application of single molecule force spectroscopy as a tool for probing structural transitions has yielded unprecedented insights into various nucleic acid structures (Chandra et al., 2017; Greenleaf et al., 2008; Mandal et al., 2019; Yang et al., 2018; Zhong et al., 2016), dynamic cellular processes (Desai et al., 2019; Wen et al., 2008) as well as mechanisms of PRF (Chen et al., 2017; Green et al., 2008; Halma et al., 2019; Yan et al., 2015).

Here we present the crystal structure of EMCV 2A revealing a novel RNA-binding fold that we term a “beta-shell”. Using a combination of biochemical and biophysical techniques, we show that 2A binds directly to the frameshift-stimulatory element in the viral RNA with nanomolar affinity and 1:1 stoichiometry, and we define the minimal RNA element required for binding. Furthermore, through site-directed mutagenesis and the use of single-molecule optical tweezers, we study the dynamics of this RNA element, both alone and in the presence of 2A. By observing short-lived intermediate states in real-time, we demonstrate that the EMCV stimulatory element exists in at least two conformations and 2A binding stabilises one of these, a putative RNA pseudoknot, increasing the force required to unwind it. Finally, we report a direct interaction of 2A with both mammalian and bacterial ribosomes. High-resolution cryo-electron microscopy (cryo-EM) characterisation of 2A in complex with initiated 70S ribosomes reveals a multivalent binding mechanism and defines the molecular basis for RNA recognition by the 2A protein. It also reveals a likely mechanism of 2A-associated translational modulation, by competing for ribosome binding with initiation factors and elongation factors. Together, our work provides a new structural framework for understanding protein-mediated frameshifting and 2A-mediated regulation of gene expression.

## Results

### Structure of EMCV 2A reveals a new RNA-binding fold

Following recombinant expression in *E. coli*, purified 2A was poorly soluble and prone to aggregation. Buffer screening by differential scanning fluorimetry (Niesen et al., 2007) indicated that the thermal stability of the protein was enhanced by salt, with high-salt buffers (~1M NaCl) greatly improving solubility. Size-exclusion chromatography coupled to multi-angle light scattering (SEC-MALS) revealed a predominantly monodisperse, monomeric sample **(Figure 1A and B**; observed mass 18032.8 Da vs 17930.34 Da calculated from the 2A sequence), with a small proportion of 2A forming dimers (observed mass 40836.0 Da). We crystallised the protein and, in the absence of a suitable molecular replacement search model, determined the structure by multiple-wavelength anomalous dispersion analysis of a selenomethionyl derivative. The asymmetric unit (ASU) of the *P*6_2_22 cell contains four copies of 2A related by non-crystallographic symmetry (NCS), and the structure was refined to 2.6 Å resolution **(Table 1)**. Unexpectedly, the four molecules are arranged as a pair of covalent ‘dimers’ with an intermolecular disulfide bond forming between surface-exposed cysteine residues (C111). This arrangement is likely an artefact of crystallisation, which took >30 days, possibly due to the gradual oxidation of C111 promoting formation of the crystalline lattice. The N-terminal 10–12 residues are disordered in all chains except B, in which they make a long-range crystal contact with a symmetry-related molecule. Similarly, C-terminal residues beyond 137 are absent or poorly ordered in all chains.

**Figure 1.**
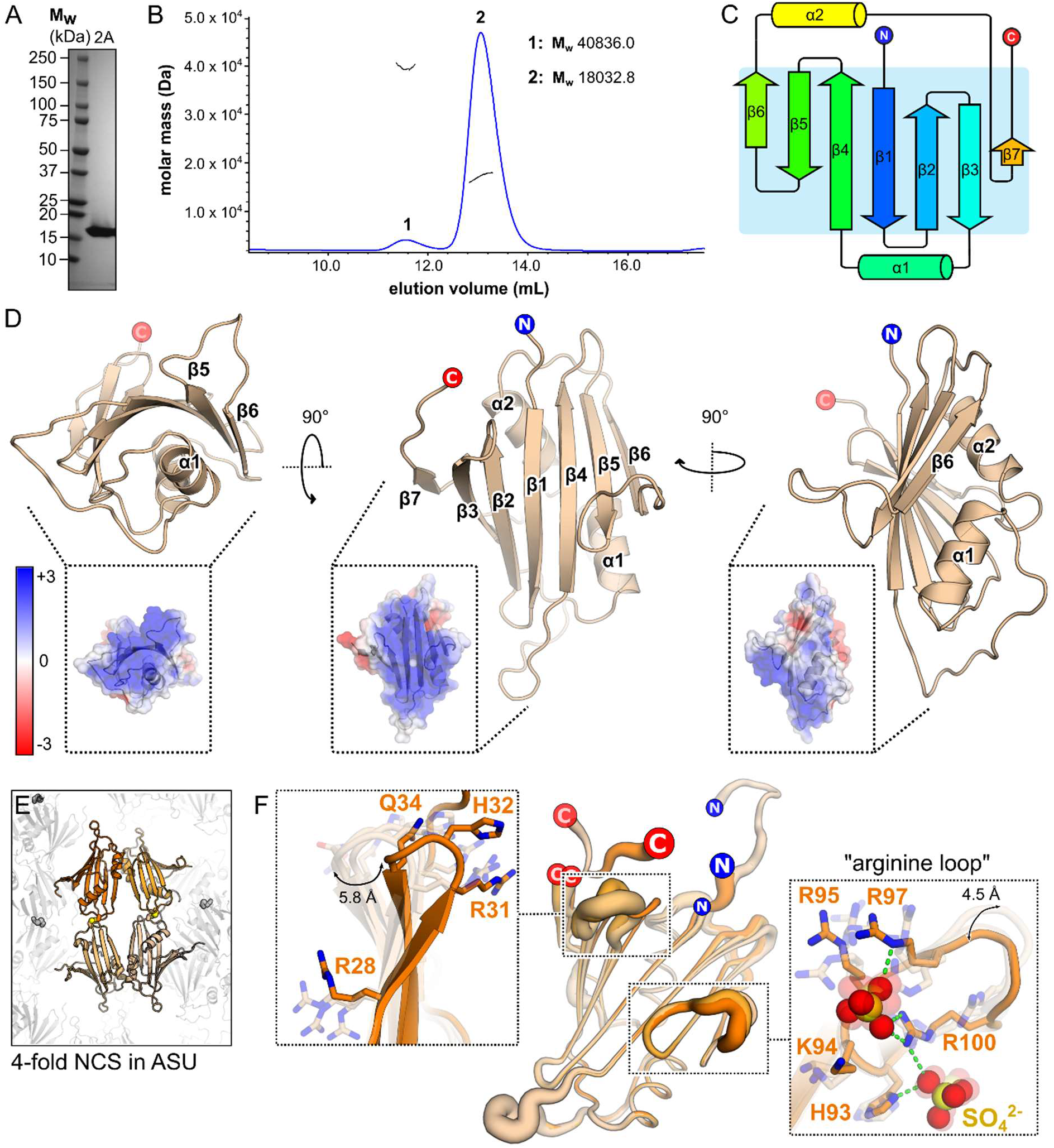
2A adopts a highly basic RNA-binding fold with intrinsic flexibility. **A)** SDS-PAGE analysis of EMCV 2A after Ni-NTA, heparin affinity and size-exclusion chromatography. The gel was stained with Coomassie blue. **B)** SEC-MALS analysis of 5.2 mg/mL 2A in high-salt buffer. The differential refractive index is shown across the elution profile (blue) and weight-averaged molar masses of the indicated peaks are listed. **C)** Topological diagram of “beta-shell” fold: a curved central sheet comprising seven antiparallel beta strands, supported by two helices. **D)** X-ray crystal structure of EMCV 2A in three orthogonal views. N- and C-termini are indicated. *<Inset>* Electrostatic surface potential calculated at pH 7.4, coloured between +3 (blue) and −3 (red) kT/e^−^. **E)** Four molecules of 2A are present in the asymmetric unit of the crystal, arranged as two pairs of disulfide-linked dimers (spheres). **F)** Superposition of the four NCS-related 2A chains in E) reveals regions of conformational flexibility. The width of the cartoon is proportional to atomic B-factor. *<Insets>* Close-up view of surface loops exhibiting the greatest variation per molecule. Flexible sidechains are shown as sticks, and the Cα backbone deviation is indicated in Å. The positions of two sulfate ions from the crystallisation buffer are indicated with spheres.

**Table 1.**
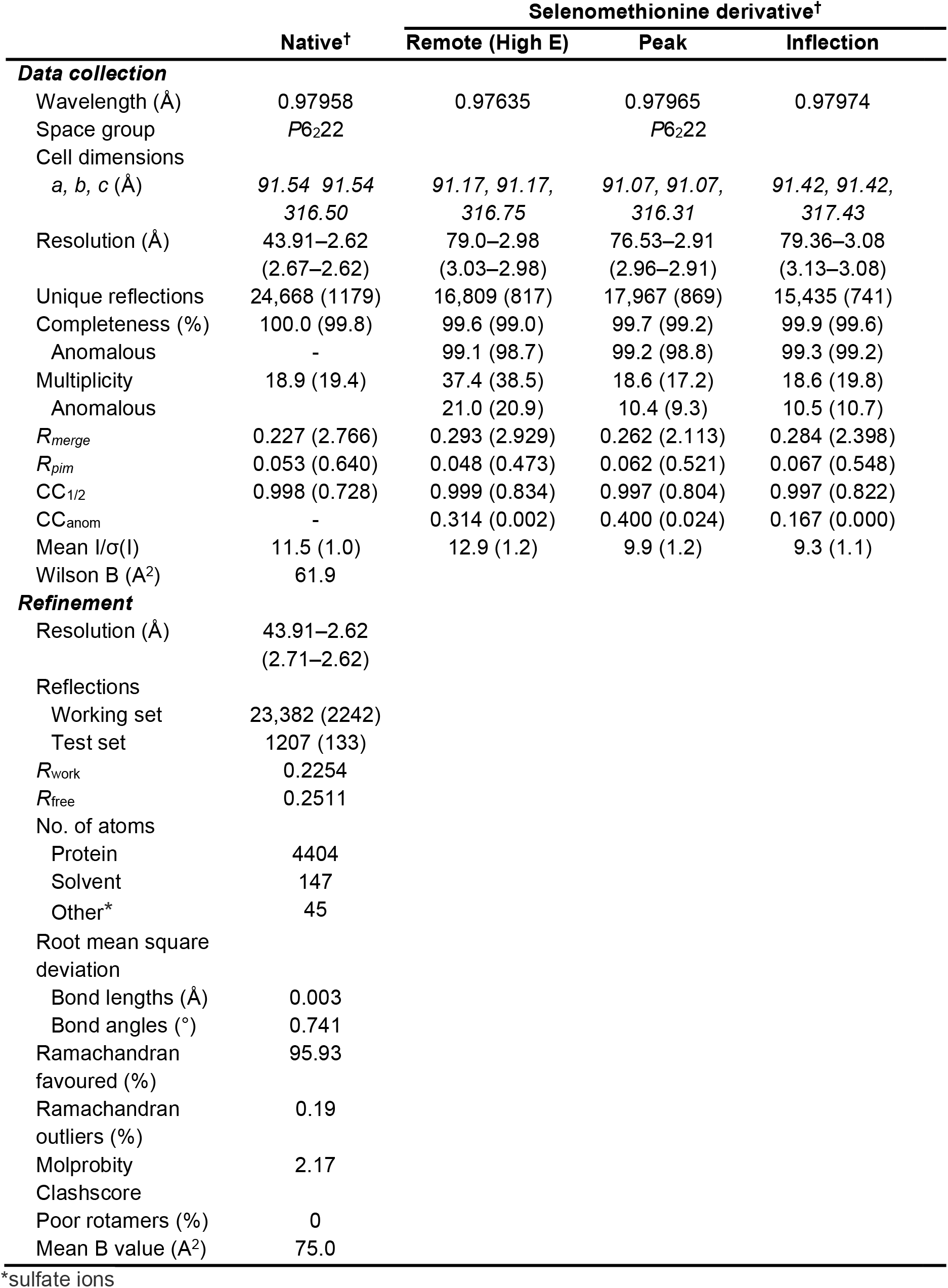
Crystallographic data collection and refinement. Data were recorded from a single native crystal and a single crystal of selenomethionine-derivatised protein. Values for high resolution shells are shown in parentheses. Related to **Figure 1**.

2A adopts a compact, globular fold of the form β_3_αβ_3_αβ **(Figure 1C)**. Given the absence of structural homology to any other protein, we term this new fold a “beta shell”. The most striking feature of this fold is a seven-stranded anti-parallel beta sheet that is highly curved **(Figure 1D)**. The concave face of the beta sheet is supported by tight packing against the two alpha helices: together, this comprises the hydrophobic core of the fold. In contrast, the solvent-exposed convex face and surrounding loops are enriched with arginine, lysine and histidine residues, conferring a strong positive electrostatic surface potential at physiological pH. This is consistent with an RNA-binding mechanism in which the negatively charged ribose phosphate backbone is recognised by electrostatic interactions (Moreira et al., 2007).

Superposition of the four NCS-related chains and an analysis of the atomic displacement factors reveals regions of flexibility within the 2A protein **(Figure 1E and F)**. In addition to the N- and C- termini, the β2-loop-β3 region (residues 28–37) exists in multiple conformations that deviate by up to 5.8 Å in the position of the C_α_ backbone. Similarly, the arginine-rich loop between β5 and β6 (“arginine loop”, residues 93–100) is highly flexible, with backbone deviations of up to 4.5 Å. Interestingly, this region has multiple roles: it acts as the 2A NLS (Groppo et al., 2011) and mutation of R95 and R97 to alanine inhibits PRF by preventing 2A binding to the stimulatory element in the mRNA (Napthine et al., 2017). In support of the latter observation, we observe that this loop binds sulfate ions (present at high concentration in the crystallisation buffer). Sulfate binding sites often indicate regions of a protein that could interact with an RNA phosphodiester backbone, based on similar geometry and charge.

Several previous studies have described mutations, truncations or deletions in EMCV 2A that affect its activity. We can now better understand the structural consequences of these alterations (Groppo et al., 2011; Petty et al., 2014; Svitkin et al., 1998). Many of the truncation mutants would lack substantial portions of secondary structure and expose elements of the 2A protein hydrophobic core **(Figure S1A and B)**. This would severely disrupt the folding of the protein and the results obtained with these mutants should be interpreted with caution. However, the loop truncation (2A_Δ94-100_) and point mutations made by Groppo et al. (Groppo *et al*., 2011) **(Figure S1C and D)** would not be predicted to disrupt the fold of 2A and can be interpreted in light of the structure. Notably, in 2A, a C-terminal YxxxxLΦ motif predicted to bind eIF4E is within a beta strand, whereas the equivalent motif in 4E-BP1 is alpha-helical (Siddiqui et al., 2012). As a result of the more extended backbone conformation in 2A, Y129 is distal to L134 and I135. It is also partially buried and anchored in place by surrounding hydrophobic residues, in contrast to the tyrosine residue in 4E-BP1 that protrudes and makes significant contacts with a pocket on the eIF4E surface. Overlay of our 2A structure with the structure of the eIF4E:4E-BP1 complex indicates that, without a significant conformational change, this motif is unlikely to represent the mechanism by which 2A recognises eIF4E **(Figure S1E)**.

### 2A binds to a minimal 47 nt pseudoknot in the viral RNA

The RNA sequence that directs PRF in EMCV consists of a G_GUU_UUU shift site, a variant of the canonical X_XXY_YYZ PRF slippery sequence, and a stimulatory stem-loop element downstream **(Figure 2A)**. The spacing between shift-site and stem-loop is 13 nt, significantly longer than that seen typically (5–9 nt) at sites of –1 PRF, and 2A protein has been proposed to bridge this gap through interaction with the stem-loop. We have previously demonstrated that three conserved cytosines in the loop are essential for 2A binding (Napthine et al., 2017) **(Figure 2A)**. To map the interaction between 2A and the stimulatory element in more detail, we prepared a series of synthetic RNAs with truncations in the shift site, loop, and 5′ and 3′ extensions on either side of the stem (EMCV 1–6; **Figure 2B)**. These were fluorescently labelled at the 5′ end, and their binding to 2A was analysed by electrophoretic mobility shift assay (EMSA; **Figure 2C)** and microscale thermophoresis (MST; **Figure 2D and Table 2)**.

**Figure 2.**
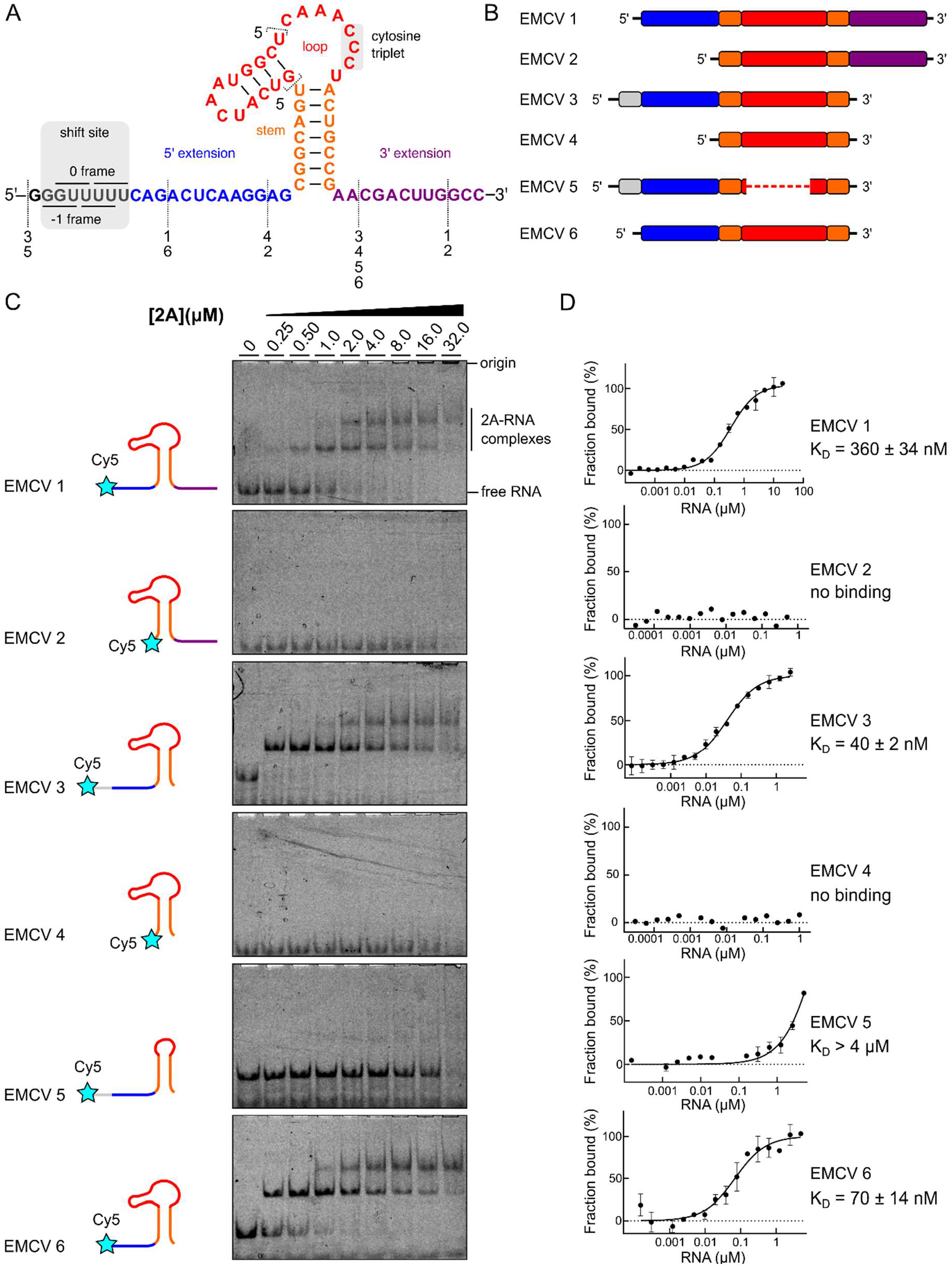
2A binds to a minimal 47 nt element in the viral RNA. **A and B)** Sequences and schematic diagrams of the EMCV 1–6 constructs used to assay 2A binding. **C)** EMSA analyses showing that removal of the 5′ extension (blue) disables 2A binding. EMSAs were conducted with Cy5-labelled EMCV RNA (50 nM) at 2A concentrations varying between 0–32 μM. Following non-denaturing electrophoresis, fluorescence was imaged using a Typhoon scanner. **D)** Microscale thermophoresis (MST) was used to quantify the interactions observed in C). Binding affinities of unlabelled 2A to fluorescently labelled EMCV RNA (5 nM) were measured using Monolith NT.Pico (NanoTemper Technologies) at 5% LED power and medium MST power. All measurements were repeated twice. RNA concentration ranges between 60 pM – 20 μM (for EMCV 1) and 150 pM – 5 μM (for EMCV 2–6).

**Table 2.** Summary of dissociation constants (K_D_) measured by microscale thermophoresis with various 2A interaction partners. Related to **Figures 2, 4 and S2**.

| | $K_D$ (LABELLED 2A) | $K_D$ (LABELLED RNA) |
| --- | --- | --- |
| **EMCV 1** | 360 ± 34 nM | 490 ± 45 nM |
| **EMCV 2** | No binding | No binding |
| **EMCV 3** | 40 ± 2 nM | 31 ± 4 nM |
| **EMCV 4** | No binding | No binding |
| **EMCV 5** | > 4 μM (does not reach saturation) | > 4 μM (does not reach saturation) |
| **EMCV 6** | 70 ± 14 nM | 50 ± 12 nM |
| **40S** | 10 ± 2 nM (apparent) | N/A |
| **60S** | > 1000 nM (apparent; does not reach saturation) | N/A |
| **30S** | 4 ± 1 nM (apparent) | N/A |
| **50S** | > 500 nM (apparent) | N/A |
| **70S** | 49 ± 13 nM (apparent) | N/A |
| **70S IC** | > 60 nM (apparent; does not reach saturation) | N/A |

Binding of 2A to EMCV 1 is high affinity (K_D_ = 360 ± 34 nM). This construct lacks the shift site, which would be within the ribosome and unavailable for 2A binding in a frameshift-relevant scenario. Removal of the 3′ extension, as in EMCV 3 and EMCV 6, further increases the affinity (KD values of 40 ± 2 and 70 ± 14 nM, respectively), perhaps by removing competing base-pairing interactions. There is no substantial difference between affinities of EMCV 3 and 6, which differ only by the presence of the shift site. Removal of the 5′ extension, as in EMCV 2 and EMCV 4, completely abolishes 2A binding, and truncation of the loop, including a putative second stem (EMCV 5) reduces binding to micromolar levels. An EMSA was also performed with an N- and C-terminally truncated version of 2A containing a C111S mutation (2A_9-136; C111S_), to probe whether the short peptide extensions added to the 2A N- and C-terminus during expression cloning or the disulfide bond observed in the crystal structure contribute to RNA binding. As seen **(Figure S2A)**, this 2A variant bound EMCV 6 RNA identically compared to the wild-type protein. Inclusion of an N-terminal Strep-II tag (SII-2A) also had no effect on RNA binding **(Figure S2A)**. In EMSAs of EMCV RNAs that bind 2A we also observe a lower-mobility species at higher protein concentrations, indicative of higher-order complex formation. To investigate the stoichiometry of binding, we performed isothermal titration calorimetry (ITC) analysis of the interaction between 2A and EMCV 6 **(Figure S2B and C)**. Although the measured K_D_ was higher (246 ± 72 nM) than observed using MST, possibly due to the higher salt concentration used to prevent 2A aggregation during the ITC experiment, the number of sites (0.87) is in good agreement with a 1:1 interaction. The largest contribution to the overall free energy of binding (ΔG, –9.02 kcal/mol) is enthalpy (ΔH, –13.9 ± 0.81 kcal/mol), consistent with an interaction mechanism driven by hydrogen bond or electrostatic contact formation. Finally, to test whether the presence of the fluorophore on the RNA affected 2A binding, we instead fluorescently labelled 2A and performed the reciprocal MST experiments with unlabelled RNA **(Figure S2D and Table 2)**. The observed K_D_ values are in good agreement between the two approaches.

To further validate these observations, we asked whether the small EMCV stem-loop RNAs could act as competitors to sequester 2A and reduce the efficiency of PRF in rabbit reticulocyte lysate (RRL) *in vitro* translation reactions programmed with an EMCV dual luciferase frameshift reporter mRNA **(Figure S2E)**. Indeed, when unlabelled EMCV 1, 3 and 6 were added in excess, they were able to compete with the stimulatory element present in the reporter, thereby reducing the amount of the –1 frame product. In contrast, EMCV 2, 4 and 5 had no such effect, reinforcing the results of direct binding experiments.

The failure of 2A to bind to EMCV 2, 4 and 5 was unexpected as these RNAs retain the main stem and the conserved cytosine triplet in the putative loop region. A possible explanation is that the frameshift-relevant state may include an interaction between the loop and the 5′ extension, forming a different conformation that 2A selectively recognises. Inspection of the primary sequences flanking the stem of the EMCV frameshift region revealed a number of possible base-pairing interactions, between 5′ or 3′ extensions and the loop, generating potential pseudoknots, and between the extensions themselves, generating an additional stem separated from the main stem by an internal loop. Whilst previous RNA structure probing data (Napthine et al., 2017) are largely consistent with the basic stem-loop model, we investigated the possibility that the EMCV PRF site forms a more complex structure by mutagenesis of the 5′ extension and loop C-triplet. Individually, G7C and C37G mutations both reduce 2A-dependent PRF to near-background levels **(Figure S3A and B)**. However, in combination, the G7C+C37G double mutation restores PRF to wild-type levels, and EMSA experiments with these mutants confirm that this is due to inhibition and restoration of 2A binding **(Figure S3C)**. Together, this demonstrates the likelihood of a base-pair between positions 7 and 37 that is necessary to form a conformation that 2A selectively recognises. Using this base pair as a restraint, RNA structure prediction (Magnus et al., 2016; Ren et al., 2005) reveals a pseudoknot-like fold **(Figure S3D)**.

### Single-molecule measurements of stimulatory element unwinding reveal multiple states

We further probed individual unfolding-refolding trajectories of the EMCV RNA variants using single-molecule optical tweezers **(Figure 3A)**. We used the force-ramp method, in which an RNA molecule hybridized to DNA:RNA handles is gradually stretched and relaxed in several cycles at a constant pulling rate. The applied force allows the RNA molecule to transition between folded and unfolded states (Chen et al., 2007; Li et al., 2006), and sudden changes in recorded force-distance trajectories are indicative of transitions between RNA conformers **(Figure 3B)**. (Chen et al., 2007; Li et al., 2006). Alongside the wild-type EMCV RNA sequence, we also tested a mutant with a substitution in the cytosine triplet (CUC), which is known to inhibit 2A binding and PRF (Napthine et al., 2017).

**Figure 3.**
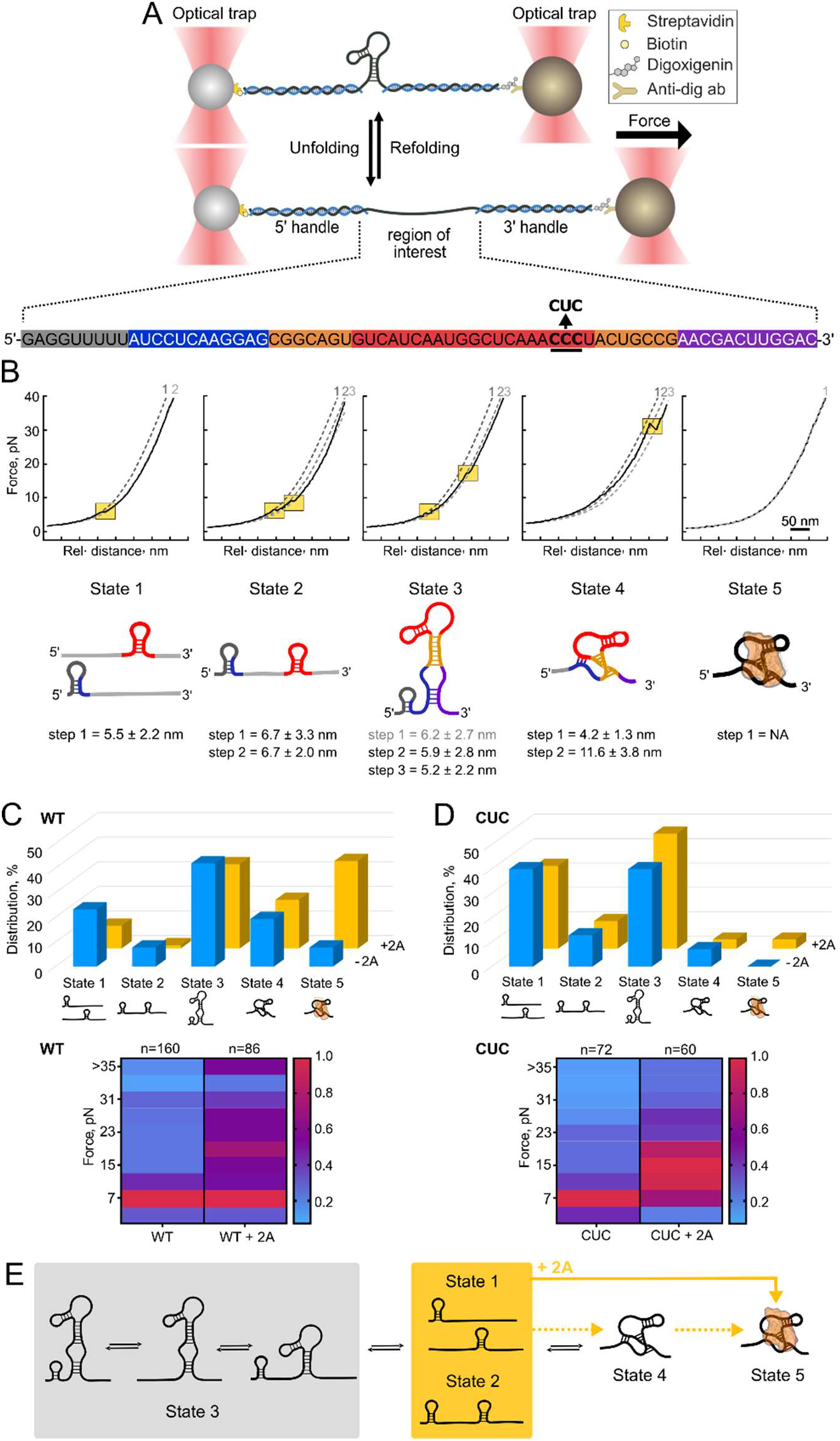
The EMCV RNA stimulatory element exists in several conformations and 2A increases its resistance to mechanical unwinding. **A)** *<Upper>* Schematic diagram illustrating the optical tweezer experiments (right). RNA is hybridized to ssDNA handles and immobilised on beads. These are used to exert pulling force on the RNA with a focused laser beam. *<Lower>* Primary sequence of the construct used in optical tweezer experiments, colour coded as in Figure 2. The location of the cytosine triplet (wild-type, WT) and point mutation (CUC) is indicated. **B)** *<Upper>* Force-distance (FD) curve examples for each of the observed states. Unfolding steps are marked with the orange squares. The numbered, dashed lines represent the fits (from left to right) of folded states, intermediates and unfolded states as described in Figure S4. *<Lower>* Inferred conformation of each state, with proposed secondary structures. Step sizes correspond to the steps observed for the WT sample. **C)** *<Upper>* Bar chart showing population distribution of RNA conformers among the measured samples. In the presence of EMCV 2A protein (WT+2A), the WT sample population profile shifts towards the more stable conformer (state 5) with a concomitant decrease in the population of the two low-force states (state 1 and 2). *<Lower>* Heatmap showing normalized distribution of unfolding forces observed in force spectroscopy experiments. We observe an increase in the higher force (~20 pN) and very high force (>35 pN) unfolding events for wild-type RNA in the presence of 2A protein (WT+2A) compared to the RNA-only sample (WT). The number of curves used for the heatmap is indicated above each column. **D)** As in C), but for CUC mutant RNA. *<Upper>* In the presence of EMCV 2A protein (CUC+2A), there is not a significant change in the population profile compared to the RNA-only sample (CUC). *<Lower>* In the presence of the 2A, (CUC+2A) there are only minor changes in the normalized distribution of unfolding forces compared to the RNA-only sample (CUC). **E)** Suggested model of the RNA conformation transitions. State 1 and 2 represent a partially folded state 3. State 3 alternates between the fully folded state (showing a three-step FD profile) and two partially folded states (showing two-step FD profiles). State 4 is a stable pseudoknot-like conformation with high unfolding forces (~30 pN). State 5 is a stable conformer population that shows no unfolding steps under the measured conditions. This state is more abundant in the presence of the EMCV 2A protein.

We initially monitored the unfolding and refolding of the wild-type (WT) and mutant (CUC) RNAs in the absence of 2A protein. The unfolding was observed in several steps representing the existence of different conformations **(Figure 3B and 3C, Table 3)**. The most dominant state (State 3) in the WT RNA was the predicted stem-loop, which was observed in 42% of molecules. This state (St3) unfolds in two steps, with the rips occurring at around 12 and 25 pN, and in some cases we observe an additional step around 6 pN. Upon release of the force, the molecule refolds with little perturbation **(Figure S4A)**. Overall, the unfolding and refolding behaviour of St3 is consistent with an extended stem-loop model with internal loop. The next population, state 4 (St4; 19%) mostly unfolds at high forces of about 30 pN, with an average step size of 11 nm **(Figure 3B)**. In some of these trajectories, we also observed a small unfolding event of 4–5 nm, occurring at low forces (8 pN). In contrast to St3, St4 showed a different refolding behaviour with a large hysteresis **(Figure S4A)**, similar to previous observations on other known pseudoknot structures (Chen et al., 2009; Ritchie et al., 2012). Refolding of the RNA occurred at lower forces of ~15 pN and in some cases, was not observed. The next state (St1; 23%) represents a population in which unfolding occurred in a single low-force step with a small extension of around 6 nm, likely characteristic of a short stem loop **(Figure 3B)**. Here, in contrast to the St3 and St4, no other unfolding step was observed up to the maximum force applied (~40 pN). In a small fraction of the traces (St2, 7%), unfolding was observed in two low-force steps of 5 and 9–10 pN, which we predict may occur if the main stem does not fold properly. Finally, 8% of the traces showed no unfolding behaviour, even above 40 pN (St5).

**Table 3.** Averaged unfolding force and extension values from the optical tweezer measurements on the wild-type (WT) and loop mutant (CUC) RNA in the presence and absence of the 2A protein. Related to **Figures 3 and S4**.

|  |  |  | **WT** | **WT + 2A** | **CUC** | **CUC + 2A** |
| --- | --- | --- | --- | --- | --- | --- |
| **State 1** | step 0 | F (pN) | 6.4±1.8 | 5.9±1.3 | 5.4±1.3 | 7.2±2.6 |
| | | $\Delta x$ (nm) | 6.2±2.7 | 5.2±1.4 | 5.4±1.7 | 5.1±1.1 |
| | step 1 | F (pN) | 10.4±4.2 | 13.5±5.2 | 9.1±4.4 | 13.3±3.7 |
| | | $\Delta x$ (nm) | 5.9±2.8 | 5.2±1.9 | 5.4±1.8 | 6.8±2.8 |
| | step 2 | F (pN) | 25.4±6.3 | 26.3±6.8 | 20.4±5.9 | 25.7±7.4 |
| | | $\Delta x$ (nm) | 5.2±2.2 | 5.4±1.6 | 5.6±2.1 | 5.1±1.9 |
| **State 2** | step 0 | F (pN) | 8.5±5.3 | | 9.7±0 | |
| | | $\Delta x$ (nm) | 4.2±1.3 | | 5.1±0 | |
| | step 1 | F (pN) | 29.2±7.9 | 30.9±6.5 | 31.1±10.6 | 21.1±0.4 |
| | | $\Delta x$ (nm) | 11.6±3.8 | 10.4±2.5 | 10.8±0.9 | 9.5±1.2 |
| **State 3** | step 0 | F (pN) | 7.0±2.3 | 8.8±3.1 | 6.4±2.7 | 13.6±4.8 |
| | | $\Delta x$ (nm) | 5.5±2.2 | 7.8±3.5 | 7.0±4.0 | 6.9±1.9 |
| **State 4** | step 0 | F (pN) | 5.3±0.9 | 5.2±0 | 5.6±1.0 | 8.7±1.8 |
| | | $\Delta x$ (nm) | 6.7±3.3 | 3.8±0 | 4.9±1.3 | 5.8±1.0 |
| | step 1 | F (pN) | 8.8±1.6 | 9.8±0 | 8.9±2.0 | 11.6±1.3 |
| | | $\Delta x$ (nm) | 6.7±2.0 | 7.0±0 | 5.6±1.9 | 8.1±1.4 |
| **State 5** | No step | F (pN) | >35 | >35 | >35 | >35 |
| | | $\Delta x$ (nm) | NA | NA | NA | NA |

Compared to the WT RNA, the main difference observed with the CUC RNA was the relative distribution of different conformers **(Figure 3D)**. The pseudoknot-like state (St4) was only seen in about 7% of the traces. This finding is consistent with our biochemical data suggesting that the cytosine triplet interacts with the 5’ extension **(Figure S3)**. Instead, St1 and St3 were observed in the majority of CUC population (both ~ 40%), and both displayed similar folding and unfolding transitions to equivalent states in WT RNA **(Figure S4A and S4B)**. This is consistent with our expectation that the predicted stem-loop would still be able to form in the CUC mutant. St2 was observed in 13% of the traces, occurring at similar forces (6 pN and 9 pN) and extensions (5 nm and 6 nm) to the WT RNA, and lastly St5 was completely absent in the CUC population.

### 2A favours the formation of an alternative state with high resistance to mechanical unwinding

We next tested how 2A binding influences RNA stability and resistance to mechanical unwinding. For the wild-type RNA, analysis of the frequency distribution of measured forces across all experiments reveals a global 2A-induced stabilisation, with increased number of unfolding events observed at high forces (~25 pN) and very high forces (>35 pN) **(Figure 3C)**. Within this population, we were able to identify the same five states based on the unfolding and extension behaviour **(Figure 3B and 3C, Table 3)**, yet the population densities showed significant differences compared to RNA-only experiments **(Figure 3C)**. The proportion of predicted stem-loop and pseudoknot-like conformations (St3 and St4, respectively) were relatively unchanged by addition of 2A, but in these populations, we no longer observed a low-force step (8 pN and 6 nm) corresponding to the unfolding of short stems immediately 5′ to the main stem loop. Strikingly, the proportion of molecules in the low-force St1 and St2 decreased in the presence of 2A (St1 23% to 9%; St2 8% to 1%), accompanied by a concomitant increase in the proportion of St5 (8% to 36%). We assume the 2A induced St5 is resistant to unwinding, and we did not observe full extension even at forces of ~40 pN **(Figure 3B and S4A)**.

Because St5 probably unfolds at forces beyond the maximum used in our experiments, we cannot determine whether the St5 conformers observed in the WT and WT+2A experiments are truly equivalent. It was also necessary to maintain a low concentration of 2A (~300 nM, close to the K_D_) to prevent aggregation and minimise non-specific interactions. Given our observed K_D_ values for 2A binding **(Figure 2D)**, it is likely that ~50% of traces in WT+2A experiments correspond to RNA-only events. In light of these caveats, several interpretations are possible. The simplest is that St5 is a distinct RNA conformation that exists in equilibrium with others and that 2A binding stabilises this conformation, thus increasing its relative abundance. Alternatively, 2A may preferentially bind to semi-folded intermediates (St1 and St2), remodelling them into a highly stable state (St5) that differs from any conformation in the absence of 2A. Conversely, St5 could simply represent a 2A-bound and stabilised version of pseudoknot St4, with the same RNA conformation and similar persistence properties, but with a higher unwinding force. In this scenario, St1 and St2 may be folding precursors of St4. Disappearance of St1 and St2 in the presence of 2A may be due to their conversion to St4 as the equilibrium shifts towards St5 formation (i.e. St1-2 ⟶ St4 ⟶ St5). In this explanation, St4 would still be observed under non-saturating 2A concentrations **(Figure 3E)**.

Finally, we tested the effects of 2A on the CUC mutant RNA. Within this population we did not observe a large, global 2A-induced stabilisation **(Figure 3D)**. Unlike for the WT RNA, the presence of 2A did not change the distribution of states **(Figure 3D and Table 3)**. In addition, St5 was largely absent, in contrast to the WT + 2A experiments in which it is the major species. The inability of the non-frameshifting CUC RNA to form St5 suggests that this may be the frameshift-relevant state: likely a stable, structured conformation that unfolds beyond the maximum forces applied (~40 pN). Fractions of St3 and St4 remained similar, and the low-force unfolding events in St2 showed a broader distribution, possibly due to non-specific interactions between 2A and the handle regions. Taken together, our results suggest that 2A binding stabilises the stimulatory RNA element and increases its resistance to mechanical unwinding.

### 2A interacts with eukaryotic and prokaryotic ribosomes

The high unwinding force of the 2A-bound St5 conformer likely reflects its role as the stimulatory element that induces a ribosomal pause at the PRF site (Napthine et al., 2019; Napthine et al., 2017). However, in addition to its role as a component of the stimulatory element, 2A has been reported to bind to 40S subunits in EMCV-infected cells (Groppo and Palmenberg, 2007). The direct interaction of 2A with ribosomes may be pertinent to its capacity to stimulate PRF: 2A may interact with translating ribosomes when they encounter the stimulatory 2A-RNA complex or (perhaps less likely) travel with elongating ribosomes to interact with the PRF signal. The 2A:40S interaction may also be relevant to the inhibition of host cell translation.

To determine if the interaction of 2A with the 40S subunit can be reproduced *ex vivo*, we purified ribosomal subunits from native RRL and analysed 2A-subunit interactions by MST **(Figures 4A and B)**. Consistent with previous data, we were unable to detect a strong interaction with 60S, but 2A forms a tight complex with 40S (apparent K_D_ = 10 ± 2 nM). To gain insight into this interaction, we prepared 2A-40S complexes for cryo-EM studies. Analysis by size-exclusion chromatography revealed that 2A co-eluted with the 40S peak, but despite extensive optimisation, subsequent cryo-EM imaging did not reveal interpretable density for 2A. As an alternative, we tested binding of 2A to purified prokaryotic ribosome subunits. 2A binds with very high affinity to 30S (apparent K_D_ = 4 ± 1 nM; **Figure 4C**), but not 50S **(Figure 4D)**. We also examined binding of 2A to intact 70S ribosomes and to reconstituted, mRNA-bound 70S ribosomes at the initiation stage (70S IC; initiator tRNA^Met^ in the P-site and an empty A-site). We were able to detect high affinity interactions with both uninitiated and initiated 70S ribosomes **(Figures 4D and E)**, implying that 50S joining and initiation complex assembly is compatible with the 2A:30S interaction.

**Figure 4.**
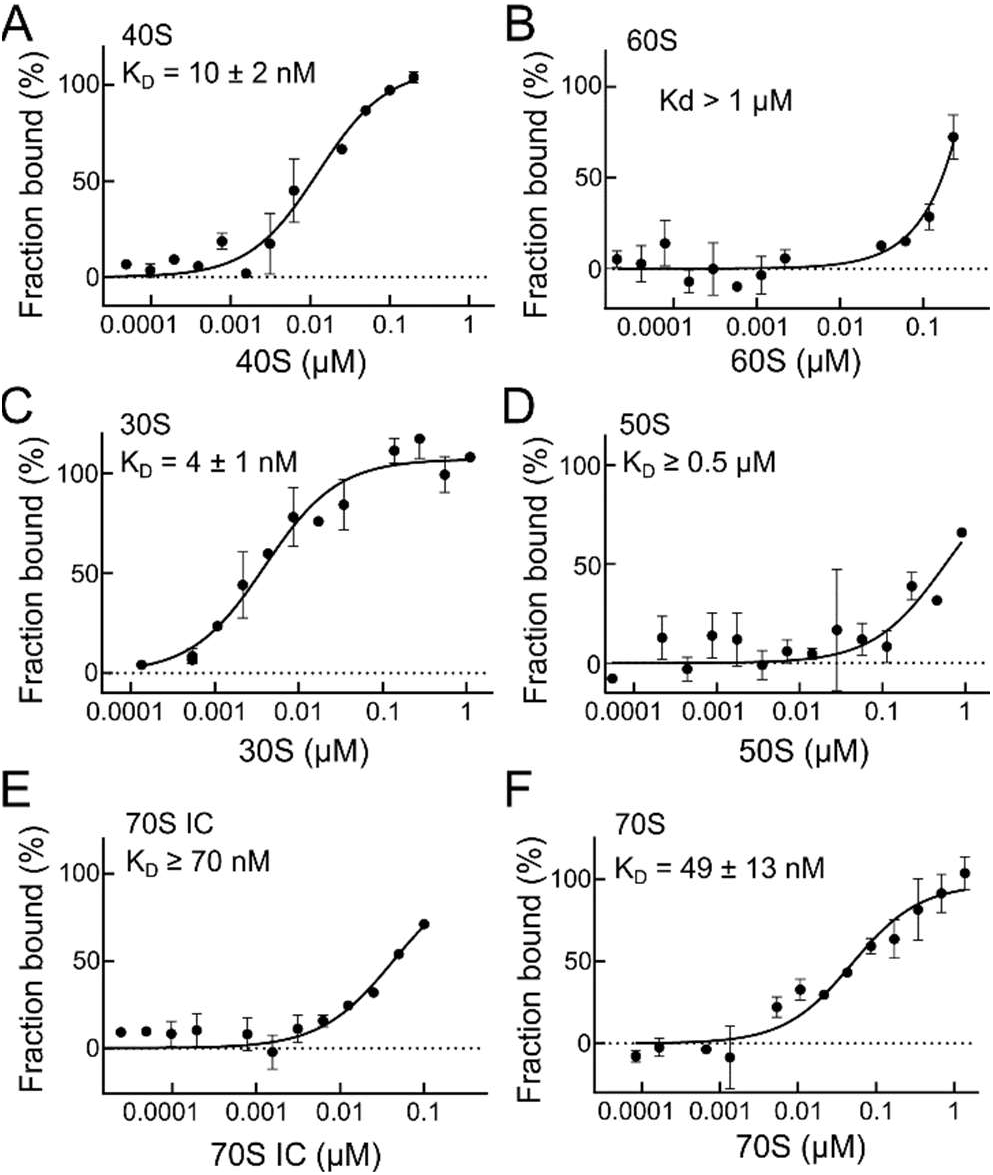
2A binds directly to eukaryotic and prokaryotic ribosomes. In all MST experiments, fluorescently labelled 2A protein was used at a final concentration of 5.0 nM. **A)** Binding curves and apparent K_D_ values using unlabelled 40S subunits at a concentration range of 20 pM – 0.4 μM. 2A binds with high affinity to the small ribosomal subunit. **B)** As in A), with 60S subunits. **C)** Binding curve and apparent K_D_ values using unlabelled 30S subunits at a concentration range of 30 pM – 1 μM. 2A shows a strong interaction with the prokaryotic small subunit. **D)** As in C), with 50S subunits at a concentration range of 27 pM – 0.9 μM. **E)** Binding curves and reported K_D_ values for 2A-70S IC interactions. **F)** Same as E, with 2A and vacant 70S.

It is well established that prokaryotic translation systems are generally responsive to eukaryotic PRF signals (Horsfield et al., 1995; Leger et al., 2004) but this has not been tested for sites of protein-dependent PRF. To address this, we measured the efficiency of the EMCV signal in a reconstituted prokaryotic translation system and in *E. coli* S30 extracts using frameshift reporter mRNAs **(Figure S5A)**. In each case, 2A-dependent PRF was observed, with ~ 15% of ribosomes changing frame. Shortening the length of the spacer to one more optimal for prokaryotic ribosomes (from 13 to 12 nt) led to a further two-fold increase in PRF. These efficiencies are comparable to those measured in eukaryotic *in vitro* translation systems (20%) (Napthine et al., 2017) and high concentrations of 2A had an inhibitory effect on translation **(Figure S5B)**, similar to that seen in eukaryotic systems.

### Cryo-EM characterisation of a 2A-ribosome complex reveals the structural basis for RNA recognition and inhibition of translation

Given the high-affinity interaction, and having validated the use of prokaryotic ribosomes as a model system to study protein-dependent PRF, we prepared complexes between 2A and the initiated 70S ribosomes and imaged them by cryo-EM **(Figure 5A and Table 4)**. After processing **(Figure S6A)**, the final 3D reconstruction produced a density map of 2.7 Å resolution **(Figure S6B–D)** and revealed three copies of 2A bound directly to 16S rRNA of the 30S subunit in a tripartite cluster **(Figure 5B and C)**. After docking the crystal structure **(Figure 1D)**, the local resolution for 2A was high enough to allow sidechain modelling and refinement **(Figure 5D)**. Alignment of the three RNA-bound conformations to the two main apo-conformations observed in different NCS-related chains of the crystal structure shows that the 2A_apo_1 (chain A-like) backbone conformation is more similar to the rRNA-bound state **(Figure S7A)**.

**Figure 5.**
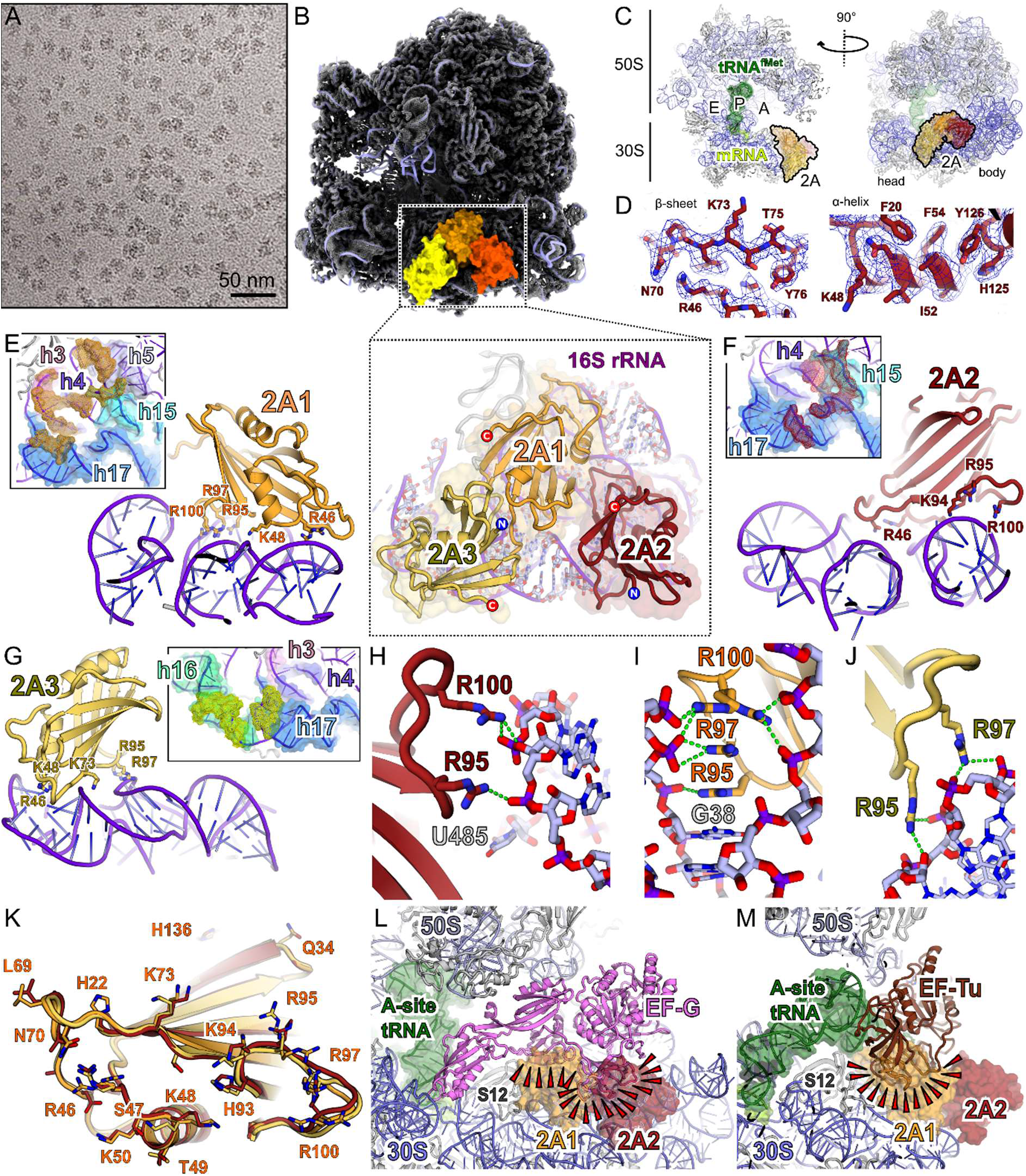
2A binds to the 70S ribosome via interactions with the 16S rRNA. **A)** Cryo-EM analysis of a complex formed between initiated *E. coli* 70S ribosomes and EMCV 2A. Images (× 75,000) were recorded on a Titan Krios microscope. **B)** Cryo-EM electron density map at 2.6 Å resolution after focus-classification and refinement. Three copies of 2A (orange, red, yellow) are bound to the 16S rRNA of the small (30S) subunit. **C)** Ribbon diagram of initiated 70S-mRNA-tRNA^fMet^-2A complex. Ribosome sites are labelled A, P and E. The initiator tRNA^fMet^ (dark green), mRNA (light green), and 2A (orange, red, yellow) are shown in two orthogonal views. **D)** Examples of local density for 2A. Well-resolved sidechains are clearly visible in both beta strands and alpha helices. E – G) Details of 2A interaction with 16S rRNA (purple). Residues involved in interactions are labelled and shown as sticks *<Inset>* View of the rRNA surface bound by each copy of 2A. The rRNA helices are colour-coded and labelled. The 2A contact surface is shown as a coloured mesh (orange, red and yellow, respectively). *H – J)* Details of interactions between 2A R95, R97 and R100 (sticks) and the rRNA backbone (sticks) for each copy of 2A (orange, red, yellow). Polar or electrostatic contacts are indicated by a green dashed line. *K)* Superposition of the three copies of 2A to highlight conformational flexibility. Residues involved in rRNA binding are labelled and shown as sticks. *L)* Comparison of 70S-2A complex to 70S pre-translocation complex with EF-G (4V7D). 2A binding would clash (red wedges) with EF-G binding. *M)* Comparison of 70S-2A complex to 70S complex with EF-Tu (5WE6). 2A binding would clash (red wedges) with EF-Tu binding.

**Table 4.**
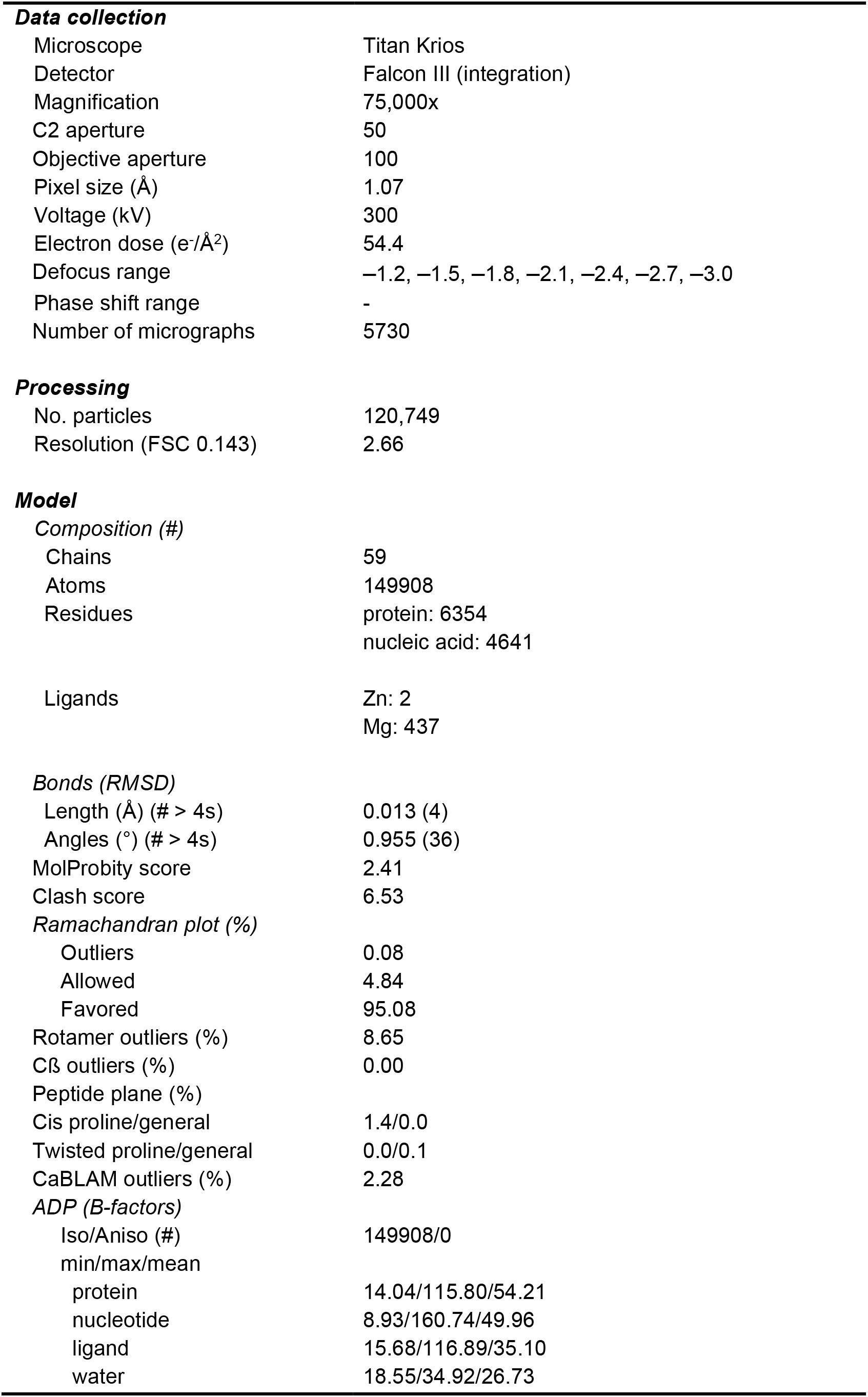
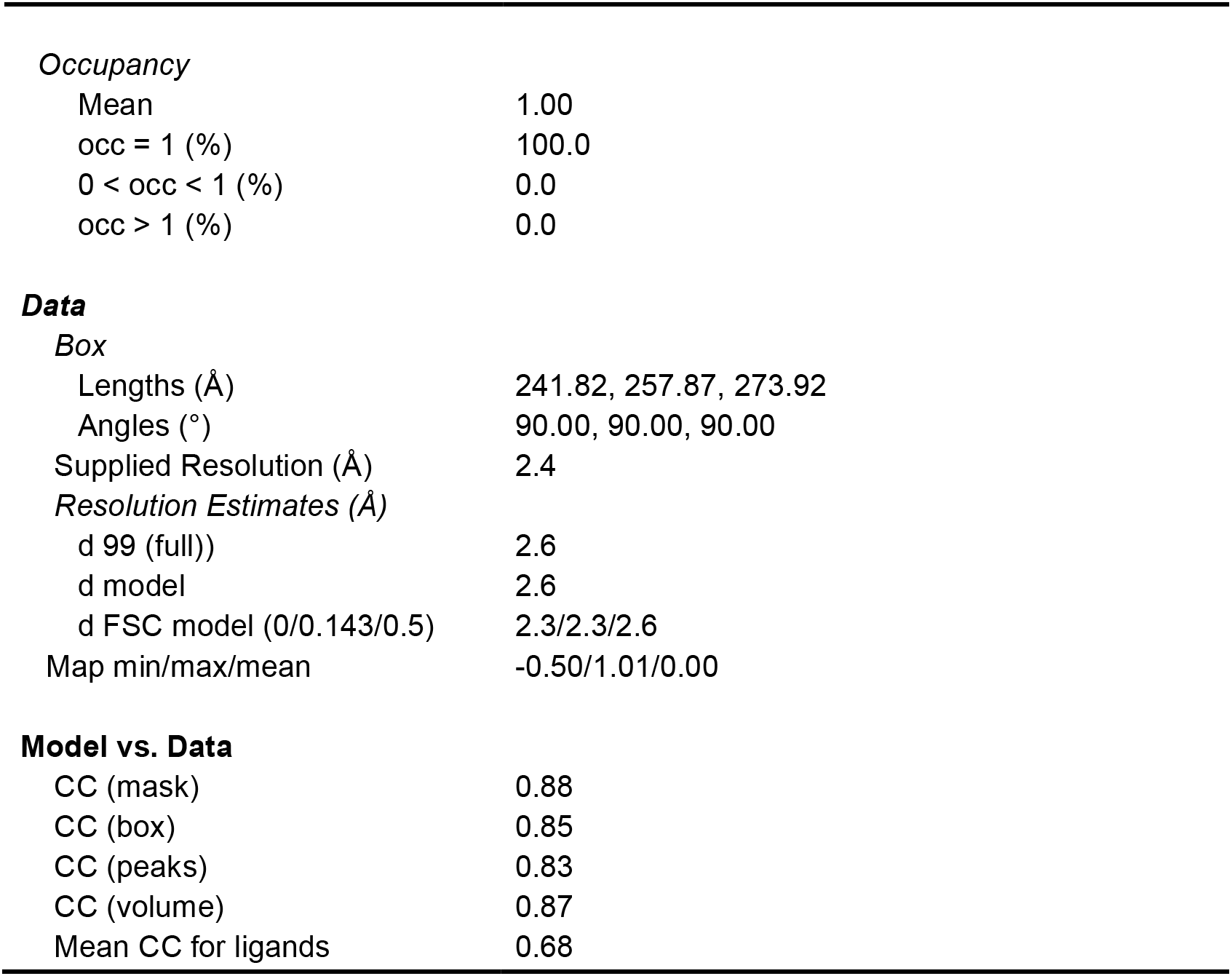
Cryo-EM data collection, processing and refinement. Related to **Figure 5, S6 and S7**.

|  |  |
| --- | --- |
| **Data collection** |  |
| Microscope | Titan Krios |
| Detector | Falcon III (integration) |
| Magnification | 75,000x |
| C2 aperture | 50 |
| Objective aperture | 100 |
| Pixel size (Å) | 1.07 |
| Voltage (kV) | 300 |
| Electron dose (e<sup>-</sup>/Å<sup>2</sup>) | 54.4 |
| Defocus range | -1.2, -1.5, -1.8, -2.1, -2.4, -2.7, -3.0 |
| Phase shift range | - |
| Number of micrographs | 5730 |
| **Processing** |  |
| No. particles | 120,749 |
| Resolution (FSC 0.143) | 2.66 |
| **Model** |  |
| *Composition (#)* |  |
| Chains | 59 |
| Atoms | 149908 |
| Residues | protein: 6354<br>nucleic acid: 4641 |
| Ligands | Zn: 2<br>Mg: 437 |
| *Bonds (RMSD)* |  |
| Length (Å) (# > 4s) | 0.013 (4) |
| Angles (°) (# > 4s) | 0.955 (36) |
| MolProbity score | 2.41 |
| Clash score | 6.53 |
| *Ramachandran plot (%)* |  |
| Outliers | 0.08 |
| Allowed | 4.84 |
| Favored | 95.08 |
| Rotamer outliers (%) | 8.65 |
| Cβ outliers (%) | 0.00 |
| Peptide plane (%) |  |
| Cis proline/general | 1.4/0.0 |
| Twisted proline/general | 0.0/0.1 |
| CaBLAM outliers (%) | 2.28 |
| *ADP (B-factors)* |  |
| Iso/Aniso (#) | 149908/0 |
| min/max/mean |  |
| protein | 14.04/115.80/54.21 |
| nucleotide | 8.93/160.74/49.96 |
| ligand | 15.68/116.89/35.10 |
| water | 18.55/34.92/26.73 |
| **Occupancy** |  |
| Mean | 1.00 |
| occ = 1 (%) | 100.0 |
| 0 < occ < 1 (%) | 0.0 |
| occ > 1 (%) | 0.0 |
| **Data****Box** |  |
| Lengths (Å) | 241.82, 257.87, 273.92 |
| Angles (°) | 90.00, 90.00, 90.00 |
| Supplied Resolution (Å) | 2.4 |
| *Resolution Estimates (Å)* |  |
| d 99 (full)) | 2.6 |
| d model | 2.6 |
| d FSC model (0/0.143/0.5) | 2.3/2.3/2.6 |
| Map min/max/mean | -0.50/1.01/0.00 |
| **Model vs. Data** |  |
| CC (mask) | 0.88 |
| CC (box) | 0.85 |
| CC (peaks) | 0.83 |
| CC (volume) | 0.87 |
| Mean CC for ligands | 0.68 |

All three copies of 2A bind directly to the ribose phosphate backbone via numerous polar and electrostatic contacts. 2A1 (orange) forms seven hydrogen bonds to rRNA helices 3, 4, 5,15 and 17, burying a surface of ~ 495 Å^2^ **(Figure 5E)**; 2A2 (red) makes 10 hydrogen bonds with helices 4, 15 and 17, burying a surface of ~ 606 Å^2^ **(Figure 5F)** and 2A3 (yellow) forms 12 hydrogen bonds the backbone of helices 16 and 17, burying a surface of ~ 532 Å^2^ **(Figure 5G)**. In all three copies of 2A, the same RNA-binding surface is involved, comprising variations of residues R46, K48, K50, K73, K94, R95 and R97. To validate this observed RNA-binding surface, we prepared variants of 2A with mutations in these residues (2A_R95A/R97A_, 2A_R46A/K48A/K50A_, 2A_K73A_; Figure S5C) and tested their ability to bind to the stimulatory element RNA **(Figure S5D)**, bind to eukaryotic ribosome subunits **(Figure S5E)** and to stimulate PRF *in vitro* **(Figure S5F)**. 2AR95A/R97A was completely functionally defective, whilst 2A_K73A_ and 2A_R46A/K48A/K50A_ exhibited moderate and mild defects in stimulation of PRF, respectively **(Figure S5G and H)**.

Interestingly, the RNA binding targets differ between the 2A binding sites. The protein does not associate with regular helices; all of the targets contain regions of helical distortion or comprise helical junctions. The most important protein residues involved in RNA binding are R95, R97 and R100, present in the flexible “arginine loop” **(Figure 1F)**. This loop adopts a different conformation in all three copies of 2A, allowing the arginine residues to bind to a wide variety of different RNA structures, not only via hydrogen bonding, but also via hydrophobic stacking interactions between exposed bases (G38) and arginine side chain guanidinium groups **(Figure 5H-J)**. Whilst base-specific contacts are rare, 2A2 interacts with U485 which is normally flipped out of helix 17 **(Figure S7B)**. Comparison of side-chain conformation at the RNA-binding surface in all three 2A molecules **(Figure 5K)** reveals a high-degree of conformational plasticity, explaining how this protein can recognise a diverse set of RNA molecules. There are also intermolecular contacts between 2A protomers. These interactions are consistent with our observations of multimers in both apo- and RNA-bound states by SEC-MALS **(Figure 1B)** and EMSA **(Figure 2C)**, and the tendency for 2A to self-associate at physiological salt concentrations. 2A1 (orange) makes hydrophobic contacts with both other molecules, burying surfaces of ~ 423 Å^2^ (2A2, red) and ~ 609 Å^2^ (2A3, yellow). There are no direct interactions between 2A2 and 2A3, and none of the observed protein-protein interfaces resemble those seen in crystal contacts. The intermolecular disulfide bond present in the crystal lattice is also absent **(Figure 1E)**.

The ribosome is in an unrotated state that would normally be elongation competent, with fMet-tRNA_i_ base-paired to the initiator codon in the P-site and mRNA available for amino-acyl tRNA delivery to the A-site (James et al., 2016). There are no 2A-induced rearrangements at the decoding centre **(Figure S7C and D)** However, the presence of 2A on the 30S subunit occludes the binding site for translational GTPases. 2A1 occupies a position that would severely clash with domain II of EF-G in both compact and extended pre- and post-translocation states (Brilot et al., 2013; Lin et al., 2015) **(Figure 5L)**. It also makes direct hydrophobic contacts with the face of S12 that would normally interact with domain III of EF-G. This 2A interaction surface on S12 is directly adjacent to the binding site for antibiotic dityromycin, which inhibits translocation by steric incompatibility with the elongated form of EF-G (Bulkley et al., 2014) **(Figure S7E)**. 2A1 would also clash significantly with domain II of EF-Tu during delivery of aminoacyl tRNAs to the A-site (Fislage et al., 2018; Loveland et al., 2020) **(Figure 5M)**. In a similar way, 2A2 would be detrimental to both EF-G and EF-Tu binding **(Figure 5L and M)**. We predict that this would have severe consequences for elongation. Indeed, at high levels, 2A is inhibitory to *in vitro* translation in both mammalian (Napthine et al., 2017) and prokaryotic systems **(Figure S5B)**. Binding at this site would also be inhibitory to initiation as it will compete for binding of IF2 during delivery of fMet-tRNA_i_ to the P-site during pre-initiation complex assembly (Hussain et al., 2016). Conversely, 2A3 occupies a site that would not clash sterically with initiation or elongation factors. Given its role in PRF, we predicted that 2A may bind proximal to the mRNA entry channel close to 30S proteins associated with mRNA unwinding activity and decoding fidelity (S3, S4 and S5) but despite extensive focussed classification, no binding at this site was observed. However, a fourth copy of 2A (2A4) was identified to bind helix 33 of the 16S rRNA ‘beak’ in the 30S head **(Figure S6C–E)**. Whilst the crystal structure could be unambiguously docked, the local resolution was insufficient for further modelling **(Figure S7E and F)**. Nevertheless, 2A4 uses a similar binding mode to recognise the distorted helical backbone.

## Discussion

Cardiovirus 2A is unique amongst picornaviral 2A proteins and a lack of homology to any known protein had precluded detailed functional inferences. Here we show that 2A adopts a novel RNA-binding fold, allowing specific recognition and stabilisation of the PRF stimulatory element in the viral RNA and direct binding to host ribosomes. The necessity for a functional Stop-Go motif at the 2A C-terminus has made a number of historical experiments difficult to interpret, as phenotypes may originate from impaired viral polyprotein processing rather than loss of specific 2A function (Groppo et al., 2011; Hahn and Palmenberg, 2001). Our structure therefore provides a framework to help rationalise several decades of preceding biochemical and virological observations.

Unusually for a multi-functional protein, it appears that many functions of 2A can be assigned to a single positively charged surface loop (residues 93–100). Despite the low pairwise sequence identity of 2A proteins amongst Cardioviruses (e.g. Theiler’s murine encephalomyelitis virus [TMEV], Saffold virus, Rat theliovirus), R95 and R97 are completely conserved. This region was originally described as an NLS as mutation of these residues, or truncation of the whole loop, abolished 2A nuclear localisation (Groppo et al., 2011). Subsequently, we demonstrated that these residues are essential for PRF activity in both EMCV and TMEV, and that their mutation to alanine prevents 2A binding to the stimulatory element (Napthine et al., 2019; Napthine et al., 2017) **(Figure S5C and E)**. Here we reveal how R95 and R97 mediate direct 2A binding to the small ribosomal subunit **(Figure 5H–J, Figure S5D)** and are therefore likely to play a critical role in conferring 2A-associated translational activities. The observed conformational heterogeneity of this loop **(Figures 1F and 5K)** indicates that mobility and flexibility are key to its myriad functions, particularly RNA binding.

Our cryo-EM structure unexpectedly revealed four distinct 2A:RNA interfaces **(Figure 5E–J and Figure S7D–E)**, providing clues as to how RNA-binding specificity is achieved. RNA recognition is driven almost exclusively by electrostatic interactions between arginine or lysine side chains and the ribose phosphate backbone oxygen atoms; very few base-specific contacts are observed. Inspection of nearby RNA chains after superposition of the three well-resolved 2A molecules failed to reveal a common preferred backbone conformation, consistent with 2A being able to flexibly bind non-regular structured RNA including features such as kinks, distortions and junctions between multiple helices. Importantly, the 70S ribosome contains many examples of A-form helices with regular geometry but 2A does not bind to any of these sites. This is consistent with our experiments to define the minimal PRF stimulatory element in the viral mRNA **(Figure 2C and D)**. Here, 2A is unable to bind EMCV 2, 4 and 5 RNAs, even though these constructs are predicted to form stable, undistorted stem-loops. Based on our biochemical data **(Figure S3)**, there is a strong likelihood that, in the 2A-bound state, the conformation of the EMCV RNA that stimulates PRF involves additional base-pairs between C-residues in the loop and a GG pair in the 5′ extension. This pseudoknot-like conformation may either pre-exist in equilibrium with other states, or it may be directly induced by 2A binding **(Figure 3E)**.

Our single-molecule data indicates that the conformational landscape of the EMCV PRF site is more complex than originally anticipated **(Figure 3B)**. Besides the predicted stem-loop (St3) and pseudoknot conformations (St4), we also observed at least two other states with low-force unfolding steps. These are likely transition intermediates—partially folded or misfolded conformations of the predicted stem-loop or pseudoknot structures. On wild-type RNA, addition of 2A reduces the prevalence of these low energy states to background levels and we see a major increase in state 5, which may represent the 2A-bound pseudoknot-like conformation. This is accompanied by a global increase in unfolding forces across the entire population **(Figure 3C)**. With CUC mutant RNA, pseudoknot-like states (St4 and St5) do not readily form and the presence of 2A induces only a slight change in unfolding trajectories **(Figure 3D)**, which may result from non-productive interactions with handle regions of the construct as observed for other nucleic acid binding proteins (McCauley et al., 2020; McCauley et al., 2015; Nir et al., 2011). Moreover, we observe minimal 2A-induced differences in the distribution of unfolding pathways **(Figure 3D)**. This supports the idea that the failure of the CUC mutant to stimulate PRF is due to its inability to adopt pseudoknot-like conformations that would normally be selectively recognised or stabilised by the 2A.

Although the 2A protein favours the stabilisation of a distinct conformer in the wild-type RNA, in our model system this state co-exists with other predicted stem-loop-like and pseudoknot-like conformations. Comparison of measured force trajectories reveals that the 2A-bound state exhibits the greatest resistance to unwinding (St5, >35 pN; **Table 3**) and therefore may cause the longest ribosomal pause, providing an extended time window for frameshifting to occur. However, given that the maximum force the ribosome can generate during translocation on an mRNA is around 13–21 pN (Liu et al., 2014), a sufficient pause is also likely to be generated by other states (St3, ~25 pN; St4 ~29 pN; **Table 3**). It was originally thought that the higher energetic stability and thus slow unfolding kinetics are important for induction of PRF (Chen et al., 2009; Zhong et al., 2016), however more recent studies report that rather than a static stability of the structure, a dynamic interplay between more conformations is crucial for efficient frameshifting (Halma et al., 2019; Ritchie et al., 2012; Ritchie et al., 2014). Thus, the observed conformational heterogeneity at the EMCV PRF site may reflect a similar requirement for stimulatory element plasticity in protein-mediated PRF.

Our current mechanistic understanding of PRF is largely informed by ensemble kinetic and single-molecule FRET studies of prokaryotic ribosomes (Bao et al., 2020; Caliskan et al., 2014; Chen et al., 2014; Choi et al., 2020; Kim et al., 2014; Qin et al., 2014). Frameshifting occurs late during the EF-G catalysed translocation step, in which the stimulatory element traps ribosomes in a rotated or hyper-rotated state, accompanied by multiple abortive EF-G binding attempts and rounds of GTP hydrolysis. A recent crystal structure showed that, in the absence of EF-G, tRNAs can spontaneously adopt a hybrid chimeric state with resultant loss of reading frame (Zhou et al., 2019). One model suggests that, in this state, an equilibrium is established between the 0 and –1 frame, which converges to 50% for long pause durations (Choi et al., 2020). Based on our structure, it is tempting to speculate that competition between EF-G and 2A binding may have a role in further prolonging the pause, thereby contributing to the high PRF efficiencies that we observe in 2A-dependent systems (Napthine et al., 2019; Napthine et al., 2017). However, the same residues in the 2A arginine loop are involved in binding both to the PRF stimulatory element in the viral RNA (Napthine et al., 2019; Napthine et al., 2017) and to ribosomal subunits **(Figure 5H–J)**. Therefore, for any given molecule of 2A, these events are likely to be mutually exclusive. This implies that the ribosome-bound form of 2A that we observe could be a secondary ‘enhancer’ of PRF efficiency, acting synergistically with the main stimulatory element. It could also be relevant to the resolution of the elongation blockade: by providing an alternate 2A-binding surface that competes with the viral RNA, the ribosome may help to induce 2A dissociation from the stimulatory element during a pause at the PRF site. Alternatively, it may not be directly relevant to frameshifting *per se*, instead representing a way of interfering with host cell translation as 2A accumulates later in infection. We cannot formally rule out the possibility that 2A3 (which does not occlude elongation factor binding, **Figure 5G**) may travel with the elongating ribosome and be ‘unloaded’ onto the PRF stimulatory element as the ribosome approaches, or may be involved in causing the ribosome to stall via protein-protein interactions between a ribosome-associated 2A and a stimulatory-element associated 2A. Indeed, the observation of a direct interaction between the ribosome and a PRF stimulatory element is not unprecedented, with a recent study revealing how the HIV-1 stem loop induces a stall by binding to the 70S A-site and preventing tRNA delivery (Bao et al., 2020). Future kinetic studies and cryo-EM imaging of ribosomes advancing codon-by-codon along the mRNA may resolve this ambiguity.

Despite our structural insights, the precise mechanism by which 2A inhibits cap-dependent initiation remains enigmatic. In normal translation, a YxxxxLΦ motif in eIF4G mediates binding to eIF4E, thereby forming eIF4F and promoting initiation. 4E-BPs also contain a YxxxxLΦ motif, competing for eIF4E binding and acting as negative regulators (Rhoads, 2009). A previous study proposed that a C-terminal YxxxxLΦ motif in 2A directly binds and sequesters eIF4E in a functionally analogous way to 4E-BP1 (Groppo et al., 2011), however our crystal structure suggests that this is unlikely to be the case without a drastic conformational rearrangement **(Figure S1E)**. It is unclear how relevant the 2A-eIF4E interaction is to host cell shut-off, as viruses harbouring mutations in the putative YxxxxLΦ motif were still able to inhibit cap-dependent translation of host mRNAs despite losing the ability to bind eIF4E (Groppo et al., 2011). An alternative explanation is that binding of 2A to 40S inhibits translation initiation. Although our cryo-EM structure suggests that 2A may block binding of IF2 to the 30S subunit, prokaryotic initiation is significantly different to cap-dependent eukaryotic initiation. Even if a similar mechanism did occur to prevent eIF2 binding in eukaryotes, this would also inhibit viral type II IRES-mediated initiation which requires all initiation factors except eIF1, eIF1A and intact eIF4F (Pestova et al., 1996a; Pestova et al., 1996b). In future it will be informative to further dissect the involvement of 2A in both IRES-dependent and cap-dependent translational initiation.

In conclusion, this work defines the structural and molecular basis for the temporally regulated ‘switch’ behind the reprogramming of viral gene expression in EMCV infection **(Figure 6)**. At the heart of this is the 2A protein: a novel RNA-binding fold with the remarkable ability to discriminate between stem-loop and pseudoknot conformers of the PRF stimulatory element. We also reveal how 2A interferes with host translation by specifically recognising distinct conformations within the ribosomal RNA. Together, this illustrates how the conformational plasticity of one RNA-binding surface can contribute to multiple functions through finely tuned relative affinities for different cellular targets.

**Figure 6.**
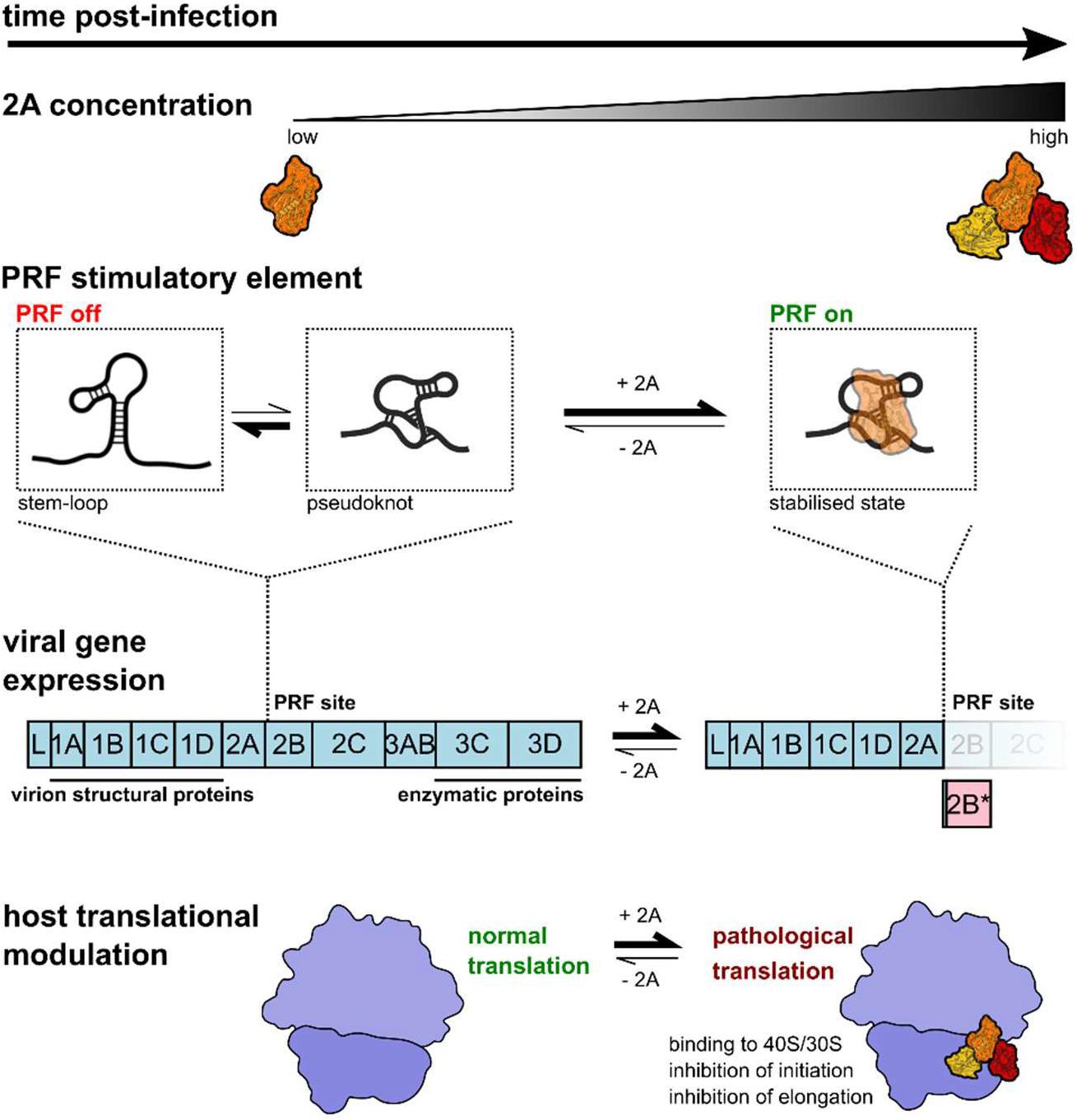
Molecular basis for 2A-induced reprogramming of gene expression. As 2A accumulates during EMCV infection, it selectively binds to and stabilises a pseudoknot-like conformation of the PRF stimulatory element, thereby enabling PRF, producing *trans*-frame product 2B* and downregulating the expression of enzymatic viral proteins later in infection. 2A also binds directly to the small ribosomal subunit at the translational GTPase factor binding site, progressively inhibiting both initiation and elongation as it accumulates. This may contribute to the shutdown of host cell translation during lytic infection.

---

**Supplementary Figure 1 - related to Figure 1.**
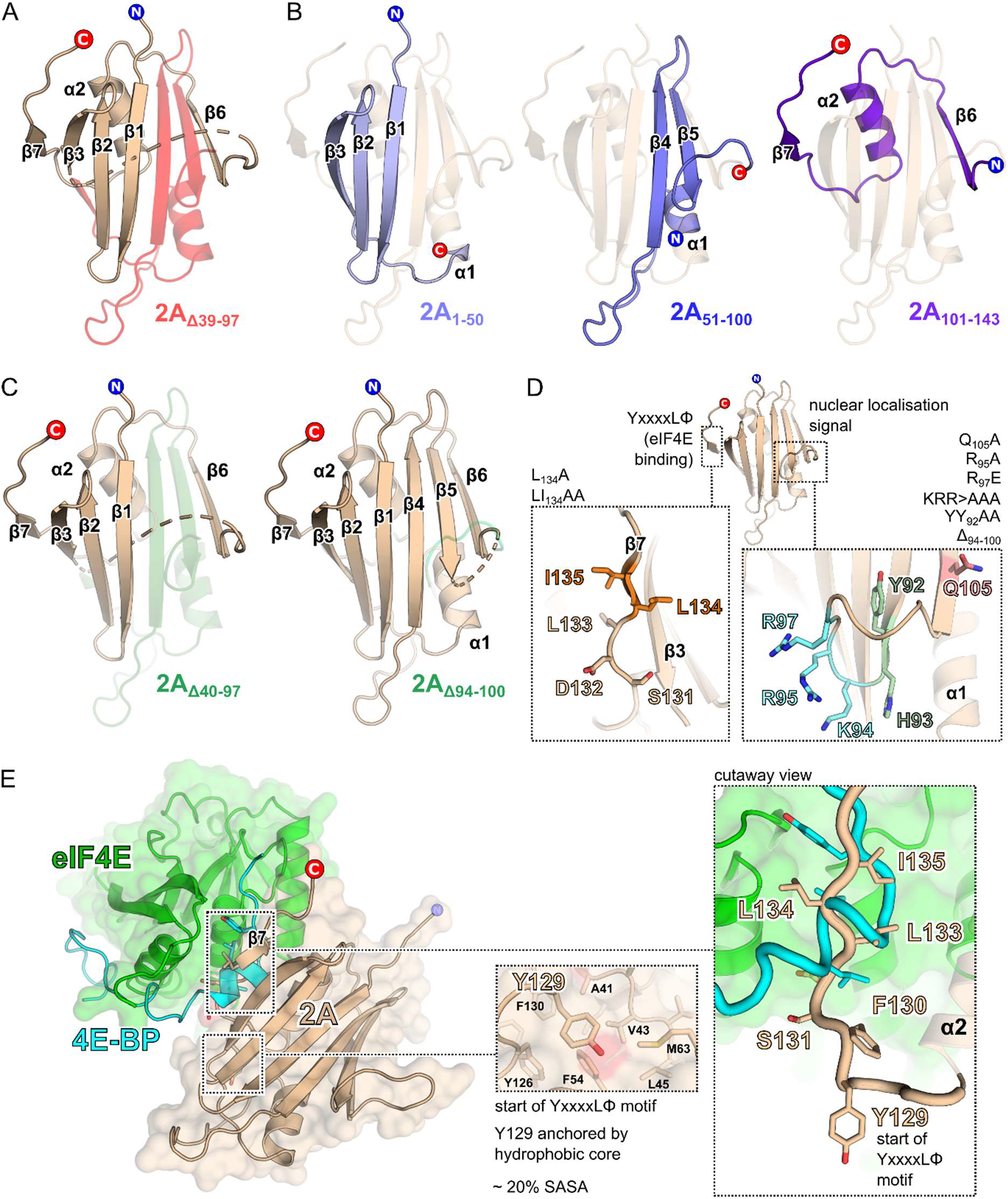
**A)** Structural consequences of the 2A_Δ39-97_ mutation described by Svitkin et al., 1998. Deleted amino acids are highlighted in red. **B)** Truncation fragments 2A_1-50_, 2A_51-100_ and 2A_101-143_ described by Petty et al., 2014. In each case the remaining fragment is highlighted in blue and overlaid against the structure of the full protein for context. **C)** Deletion mutants 2A_Δ40-97_ and 2_AΔ94-100_ as described by Groppo et al., 2011. Deleted amino acids are highlighted in green. **D)** Location of point-mutations made by Groppo *et al*. in the putative nuclear localisation sequence and putative C-terminal YxxxxLΦ eIF4E binding motif. Mutated amino acids are shown as coloured sticks. **E)** Comparison of 4E-BP1 YxxxxLΦ binding motif and the putative YxxxxLΦ motif in 2A. The crystal structure of the complex between eIF4E (green) and 4E-BP1 (blue) is shown (Siddiqui et al., 3U7X) with 2A (wheat) docked via least-squares superposition of the YxxxxLΦ motif. *<Insets>* Contrast between the 2A YxxxxLΦ motif, in an extended β-strand conformation (wheat), and the 4E-BP1 YxxxxLΦ motif, in a compact α-helical conformation, with Y129 partially buried (~ 20% solvent-accessible surface area; SASA). 2A binding to eIF4E s thus not compatible with the known 4E-BP1 interface without substantial conformational rearrangement.

**Supplementary Figure 2 - related to Figure 2.**
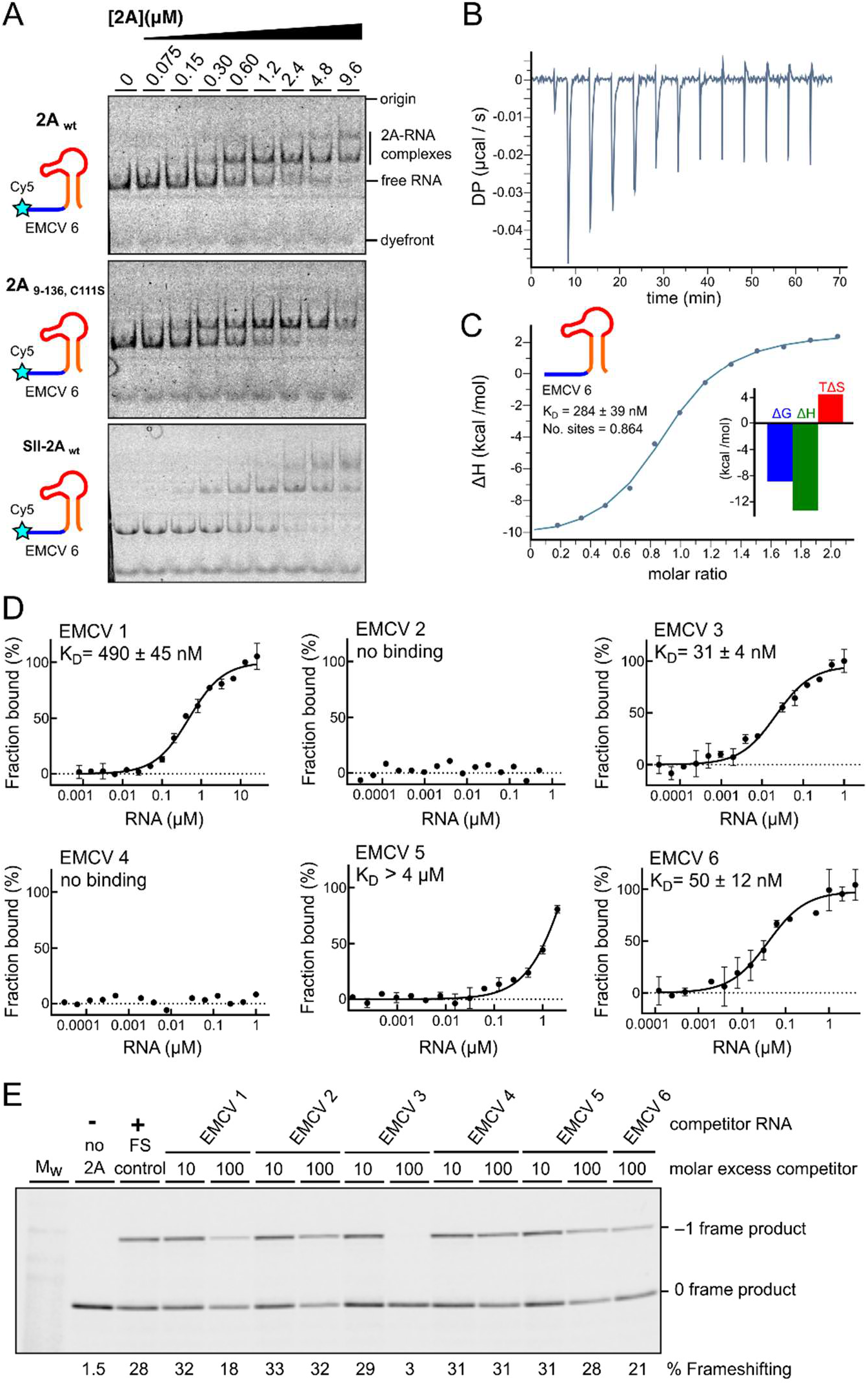
**A)** Side-by-side comparisons of 2A_wt_, 2A_9-136_; C111S and SII-2A_wt_. Equivalent RNA binding is observed in all cases by EMSA analyses conducted with 50 nM Cy5-labelled EMCV 6 RNA and 2A concentrations between zero and 9.6 μM. Following non-denaturing electrophoresis, fluorescence was imaged using a Typhoon scanner. **B)** Baseline-corrected differential power (DP) versus time for ITC titration of EMCV 6 RNA into 2A protein. **C)** Normalized binding curve showing integrated changes in enthalpy (ΔH) against molar ratio for titration in B), showing a ~1:1 molar ratio and nanomolar affinity *<Inset>* Histogram showing relative contributions of ΔH and TΔS terms to the overall exergonic interaction. **D)** MST binding curves and reported K_D_ values of fluorescently labelled 2A protein (5 nM) and short unlabelled RNAs (as in Figure 2A and B) at concentrations between 800 pM – 26 μM for EMCV 1 and 120 pM – 4 μM for EMCV 2–6. **E)** Experiment showing the effects of titrating excess short RNAs (TMEV 1–6) as competitors into an *in vitro* frameshift reporter assay. The concentrations of the reporter mRNA and 2A were kept constant in the RRL and short RNAs were added in 10- and 100-fold molar excess relative to the reporter mRNA, as indicated. Translation products were visualised by using ^35^S-Met autoradiography, and % frameshifting was calculated following densitometry and correction for the number of methionines present in 0 frame and –1 frame products.

**Supplementary Figure 3 - related to Figure 2.**
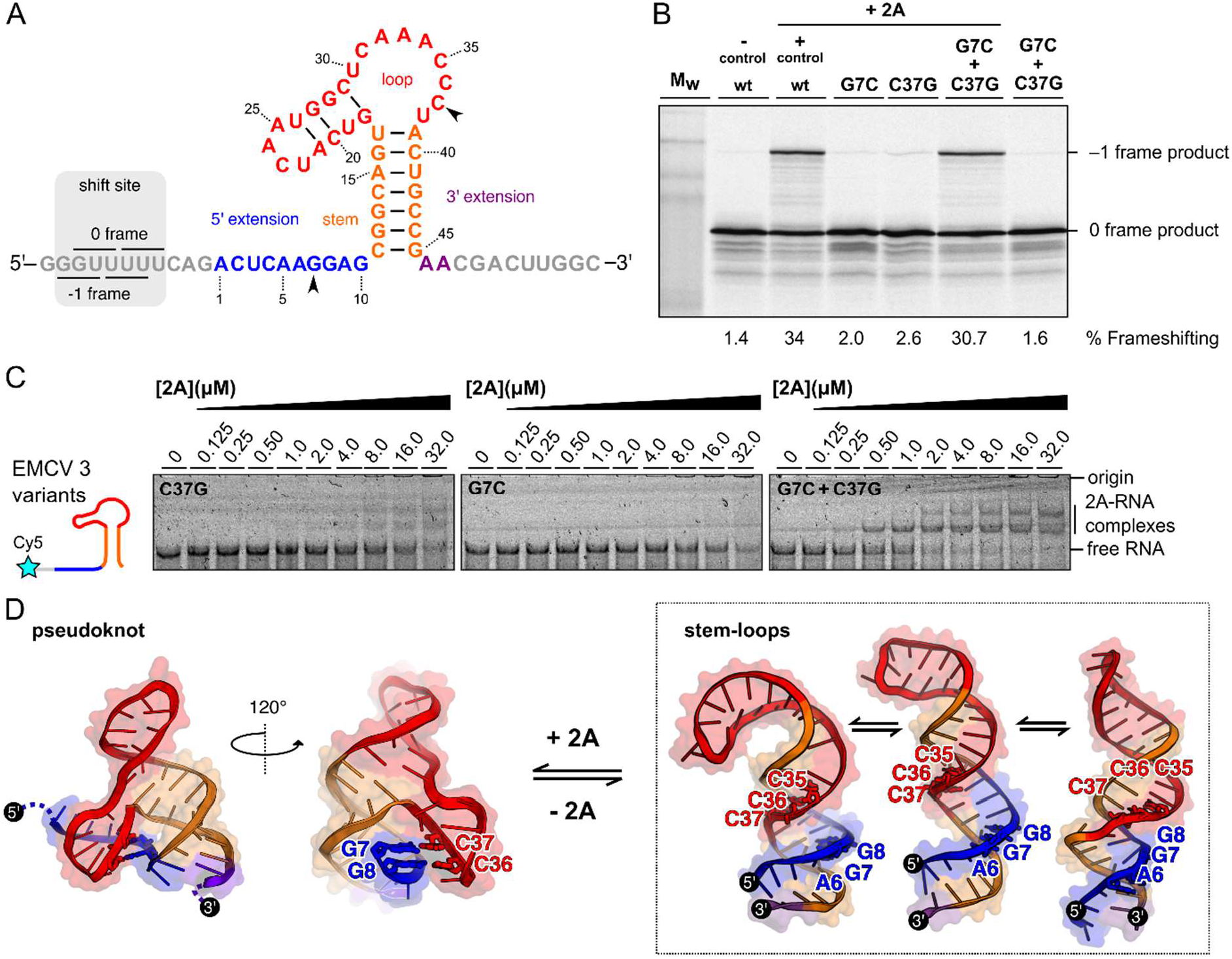
**A)** Schematic diagram showing numbered sequence of the EMCV 6 minimal PRF stimulatory element. **B)** Frameshifting assays showing evidence for a base-pairing interaction between G7 and C37. Individual G7C and C37C mutations reduce frameshifting to near-background levels. However, the double mutation (which would permit a compensatory C-G base-pair to form) restores frameshifting to wild-type levels. **C)** EMSA analyses showing that individual G7C and C37G mutations in the EMCV 6 RNA prevent 2A binding, but the double G7C+C37G mutation restores binding. Experiments were conducted with 50 nM Cy5-labelled EMCV 3 RNA variants and 2A concentrations between zero and 32 μM. Following non-denaturing electrophoresis, fluorescence was imaged using a Typhoon scanner. **D)** Equilibrium between several predicted stem-loops and alternate pseudoknot conformation, colour-coded as in A). Pseudoknot-like conformation involves base pairs between G7 and G8 in the 5′ extension and C36 and C37 in the loop (shown as sticks). These interactions are not maintained in any predicted stem-loop conformation.

**Supplementary Figure 4 - related to Figure 3.**
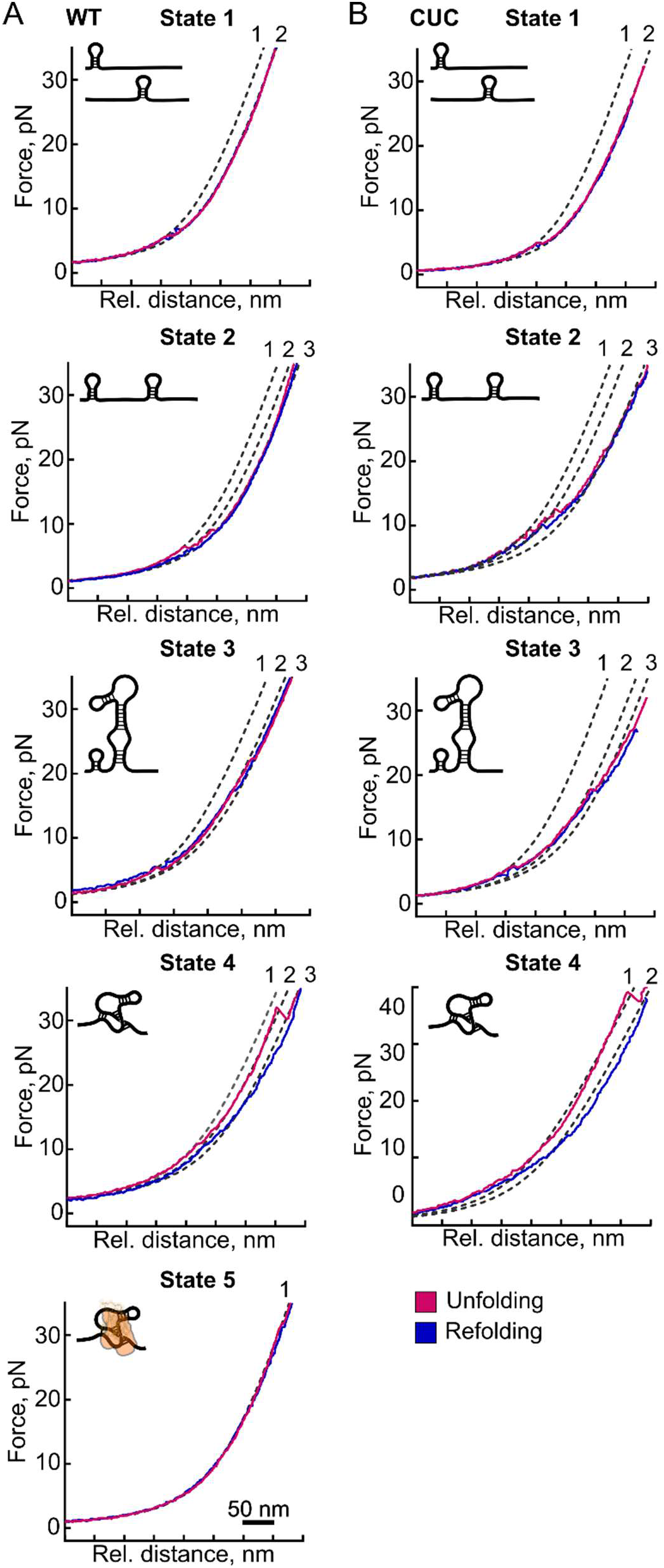
**A)** Representative force-distance curves of the unfolding (pink) and refolding (blue) transitions of the wild type (WT) CCC RNA element. An example is shown for each of the five states, and for each state, inferred conformations are depicted in the upper-left corner of each graph. The observed (un)folding steps are indicated by the dashed lines with numbers 1, 2 and 3, corresponding to folded, intermediate and unfolded states, respectively. **B)** As in A), but for the CUC mutant RNA.

**Supplementary Figure 5 - related to Figure 5.**
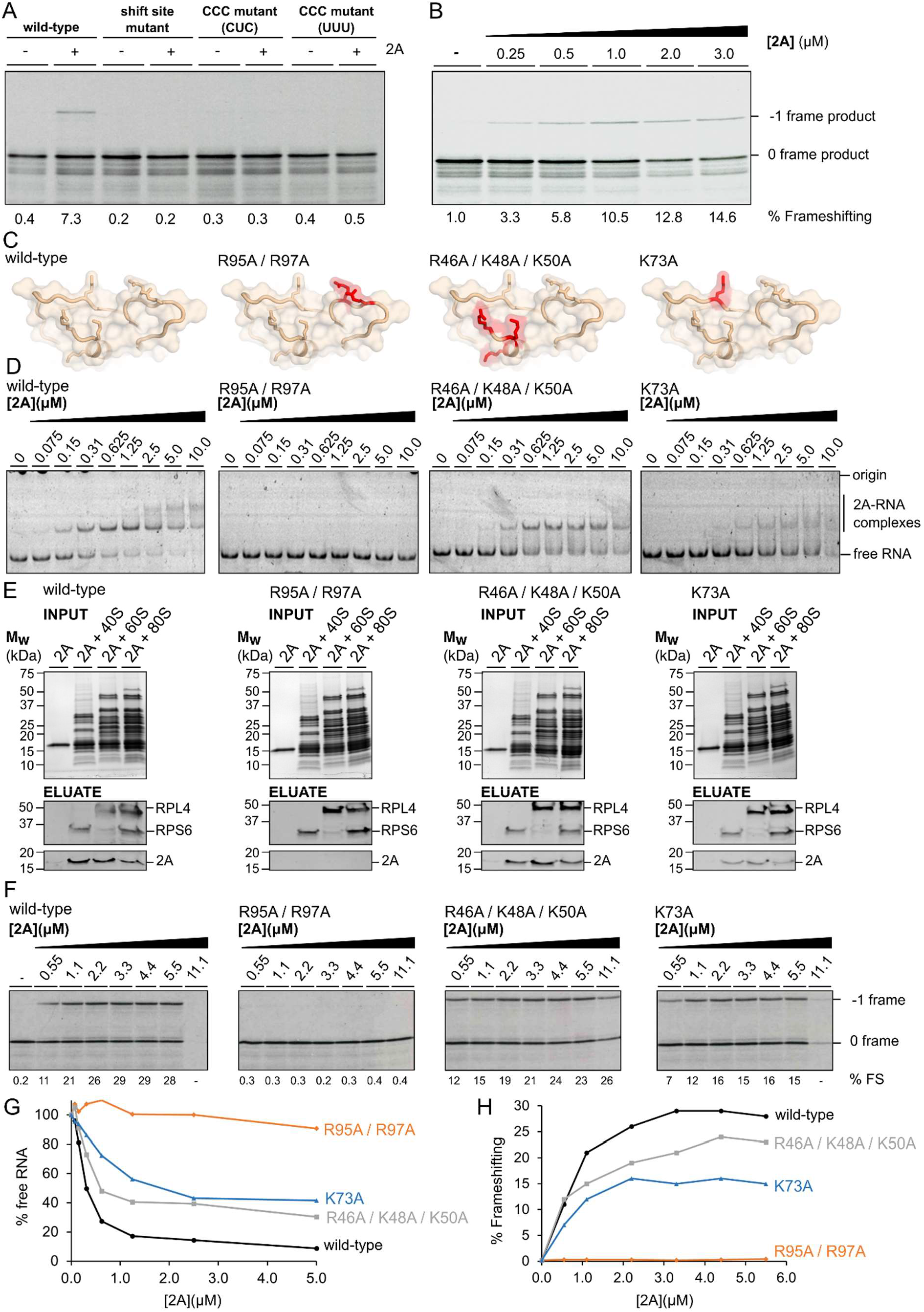
**A)** Frameshifting assay showing the reconstitution of 2A-dependent PRF in a prokaryotic *in vitro* translation system. Translation products were visualised by ^35^S-Met autoradiography and % frameshifting was calculated following densitometry and correction for the number of methionines present in 0 frame and – 1 frame products. **B)** PRF efficiency in the prokaryotic system is proportional to 2A concentration. At high levels, 2A displays inhibitory effects on total translation. Data analysed as above. **C)** Mutagenesis of residues at the 2A RNA binding surface observed in the 70S_IC_-2A structure. The locations of mutations R95A/R97A, R46A/K48A/K50A and K73A are highlighted in red and shown as sticks. **D)** EMSA analyses showing effects of the above mutations on stimulatory element RNA binding, compared to a wild-type control. Panels ordered left-right, as in A). Experiments were conducted with 50 nM Cy5-labelled EMCV 6 RNA and 2A concentrations between zero and 10 μM. Following non-denaturing electrophoresis, fluorescence was imaged using a Typhoon scanner. **E)** Gel-filtration experiments showing effects of the above mutations on eukaryotic ribosome binding, compared to a wild-type control. Panels ordered left-right, as in A). Excess 2A (2.5 μM) was incubated with 40S, 60S and 80S ribosomes (0.4 μM) prior to gel filtration chromatography using S200HR spin columns. In these experiments, 2A will only elute if bound to ribosomes. The input *<Upper>* was analysed by 4-20% gradient SDS page and visualised by staining with Imperial protein stain. The eluate *<Lower>* was analysed by western blot to detect 2A, RPS6 (small ribosomal subunit) and RPL4 (large ribosomal subunit). **F)** Frameshift reporter assays showing the effects of the above mutations on *in vitro* translation in RRL. Translation products were visualised by ^35^S-Met autoradiography and % frameshifting was calculated following densitometry and correction for the number of methionines present in 0 frame and –1 frame products. *G)* Plot of % Free RNA vs. 2A concentration, based on densitometric quantification of EMSAs in B). *H)* Plot of % Frameshifting vs. 2A concentration, based on quantification reported in D).

**Supplementary Figure 6 - related to Figure 5.**
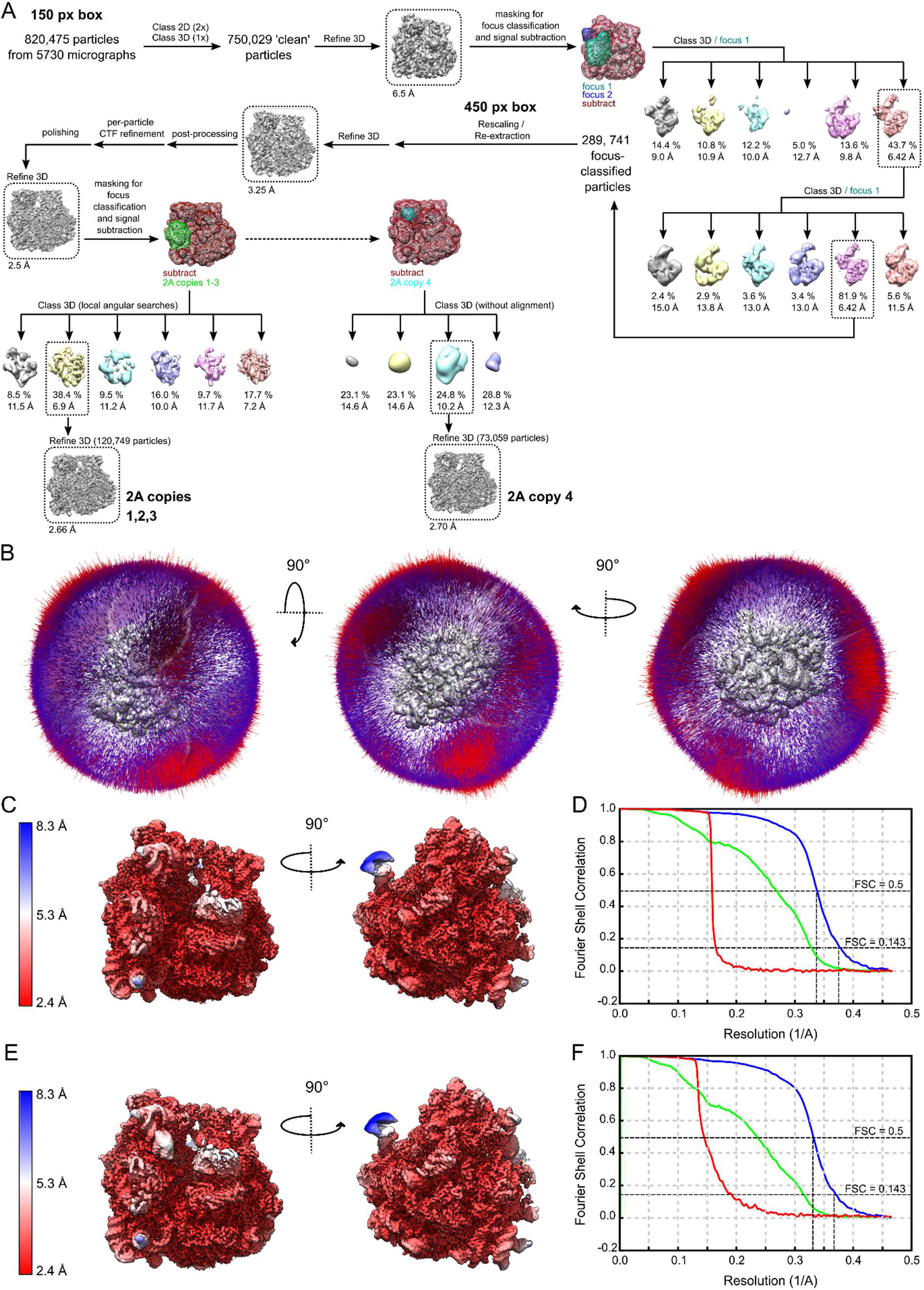
**A)** Schematic summary of steps in cryo-EM data processing. **B)** Three orthogonal views showing the angular distribution of particles contributing to the final 3D reconstruction. This is shown for the highest-resolution Refine3D result i.e. immediately after particle polishing. **C)** Local-resolution map for the final reconstruction of 70S-2A_3_. The surface is coloured by local resolution from red (highest; 2.4 Å) to blue (lowest; 8.3 Å). **D)** Gold-standard Fourier shell correlation (FSC) curve for the 70S-2A_3_ map. Masked (blue), unmasked (green) and phase-randomised masked (red) plots are shown. **E)** Local-resolution map for the final reconstruction of 70S-2A_4_, details as in C). Local resolution estimate for 2A4 is ~ 5 – 7 Å. **F)** Gold-standard Fourier shell correlation (FSC) curve for the 70S-2A_4_ map. Details as in D).

**Supplementary Figure 7 - related to Figure 5.**
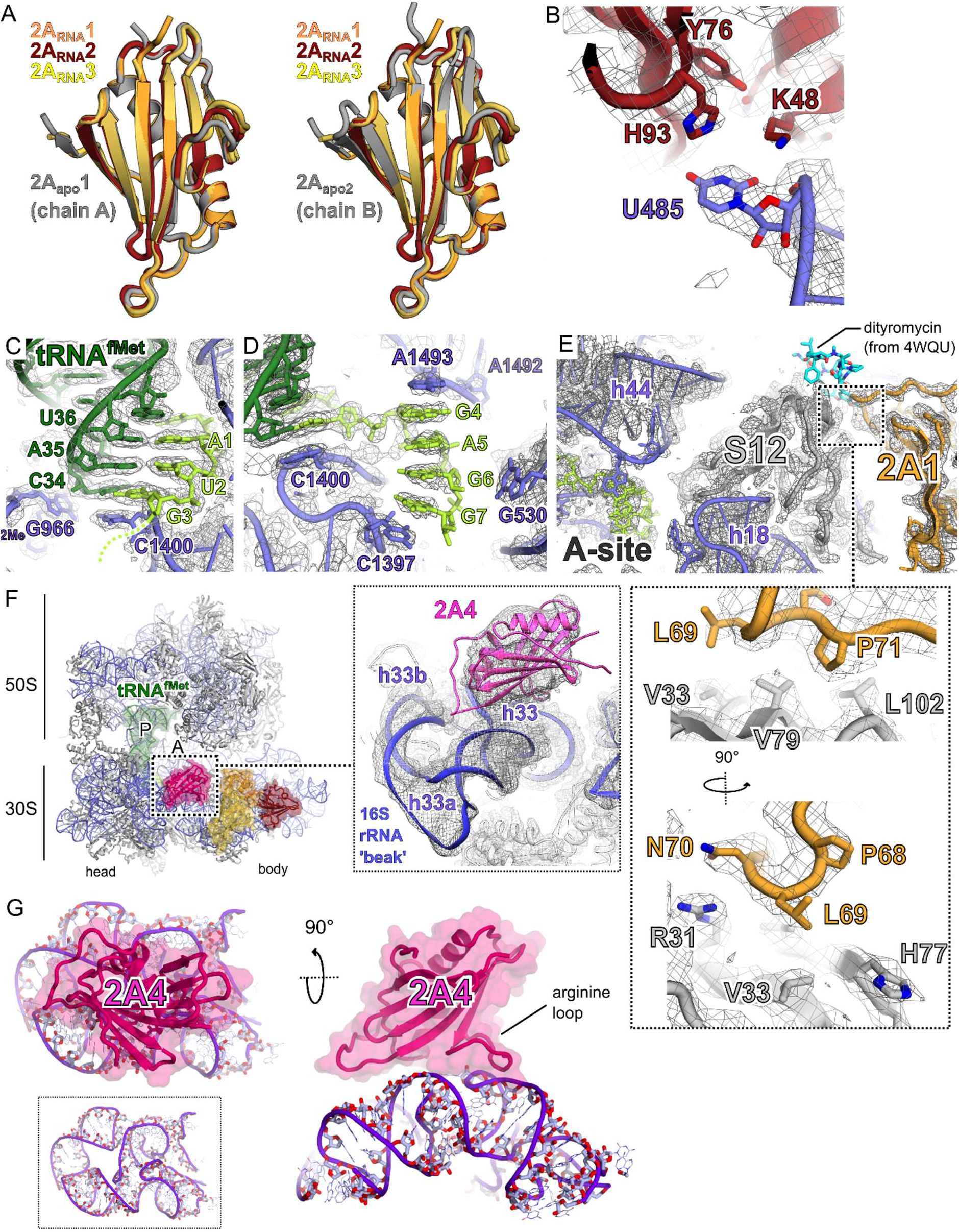
**A)** Comparison between conformations of 2A protein in RNA bound states (orange, red, yellow) and the two unliganded states observed by NCS in the crystal structure. The 2A_apo_1 conformation observed in chain A is most similar to the RNA-bound state. Structural alignments were performed by least-squares superposition of the C_α_ backbone. **B)** Details of a base-specific interaction between U485 (helix 17 of 16S) and a pocket on the surface of 2A2 (red). **C)** Cryo-EM density at the P-site. Codon-anticodon pairing between the mRNA (lime) and the initiator tRNA^fMet^ (dark green). The tRNA is in an undistorted P/P conformation as expected. **D)** Cryo-EM density at the A-site, coloured as in B). Additional 30S residues with roles in decoding are shown as sticks (purple). **E)** Details of a hydrophobic 2A1 interaction with ribosomal protein S12. The contact surface is on the factor-binding face of S12, away from the decoding centre. The binding site of antibiotic dityromycin on S12 (from 4WQU) is shown with blue sticks. **F)** Ribbon diagram of initiated 70S-mRNA-tRNA^fMet^-2A complex showing the location of the fourth copy of 2A (pink) present in a smaller population of particles. Ribosome sites are labelled A and P. The initiator tRNA^fMet^ (dark green), mRNA (lime), 2A1 (orange), 2A2 (red) and 2A3 (yellow) are also shown. *<Inset>* Section of the 70S-2A_4_ local resolution map showing electron density at the 2A4 binding site. 2A4 binds to the 3′ major ‘beak’ domain of the 16S rRNA present in the 30S ‘head’, via electrostatic interactions with the ribose phosphate backbone of helices 33 and 34. **G)** Details of 2A4 interaction with 16S rRNA (purple) in two orthogonal views.

## Materials and Methods

### Resource Availability

#### Lead contact

Further information and requests for resources should be directed to and will be fulfilled by the Lead Contact, Ian Brierley.

#### Materials availability

- Plasmids generated in this study (**see Key Resources Table**) are available on request from the lead contact.
- Purified proteins generated in this study (**see Key Resources Table**) are available on request from the lead contact.
- DNA and RNA oligonucleotides are standard synthetic products that are commercially available (see **Table 5**).

**Table 5.**
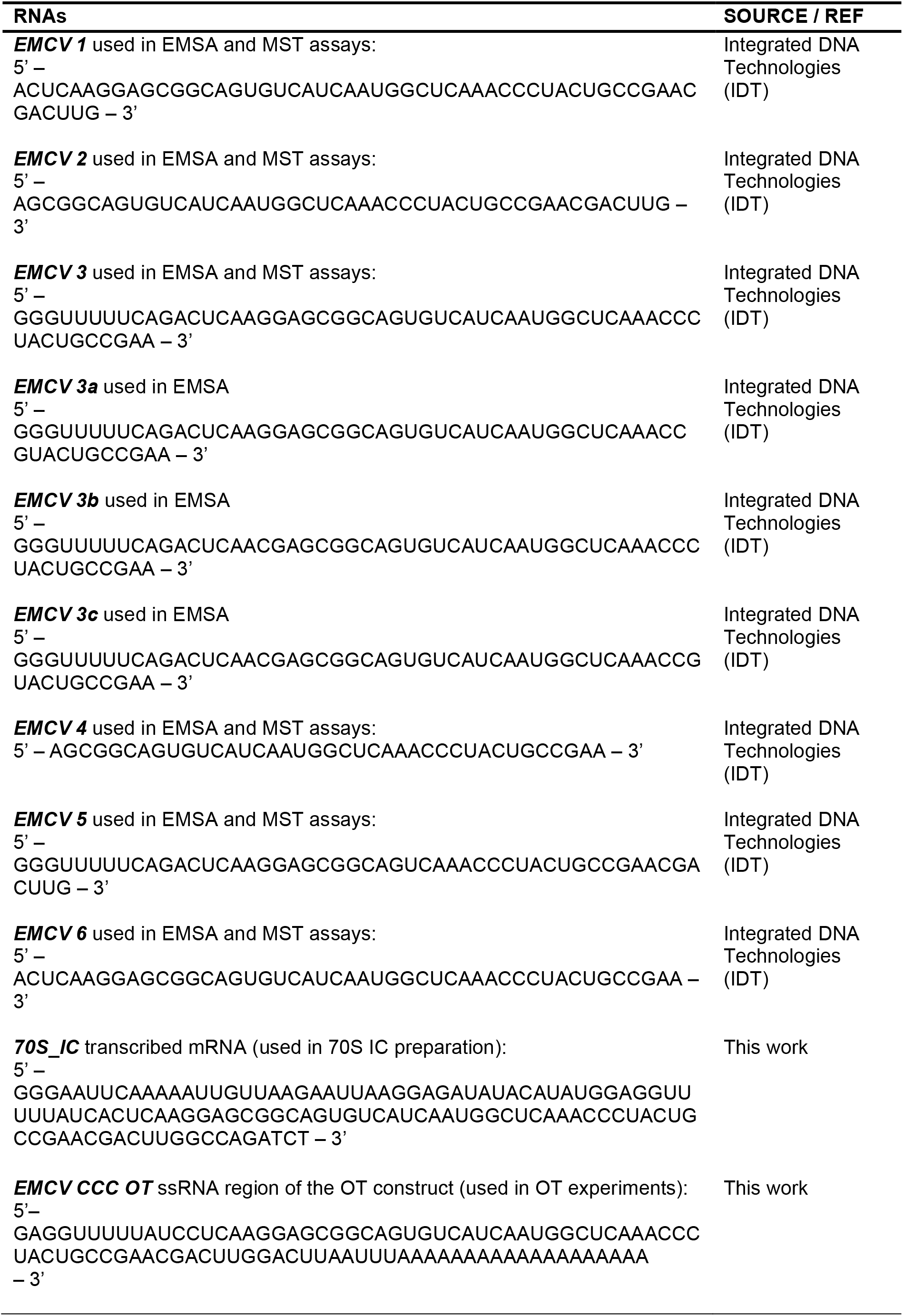
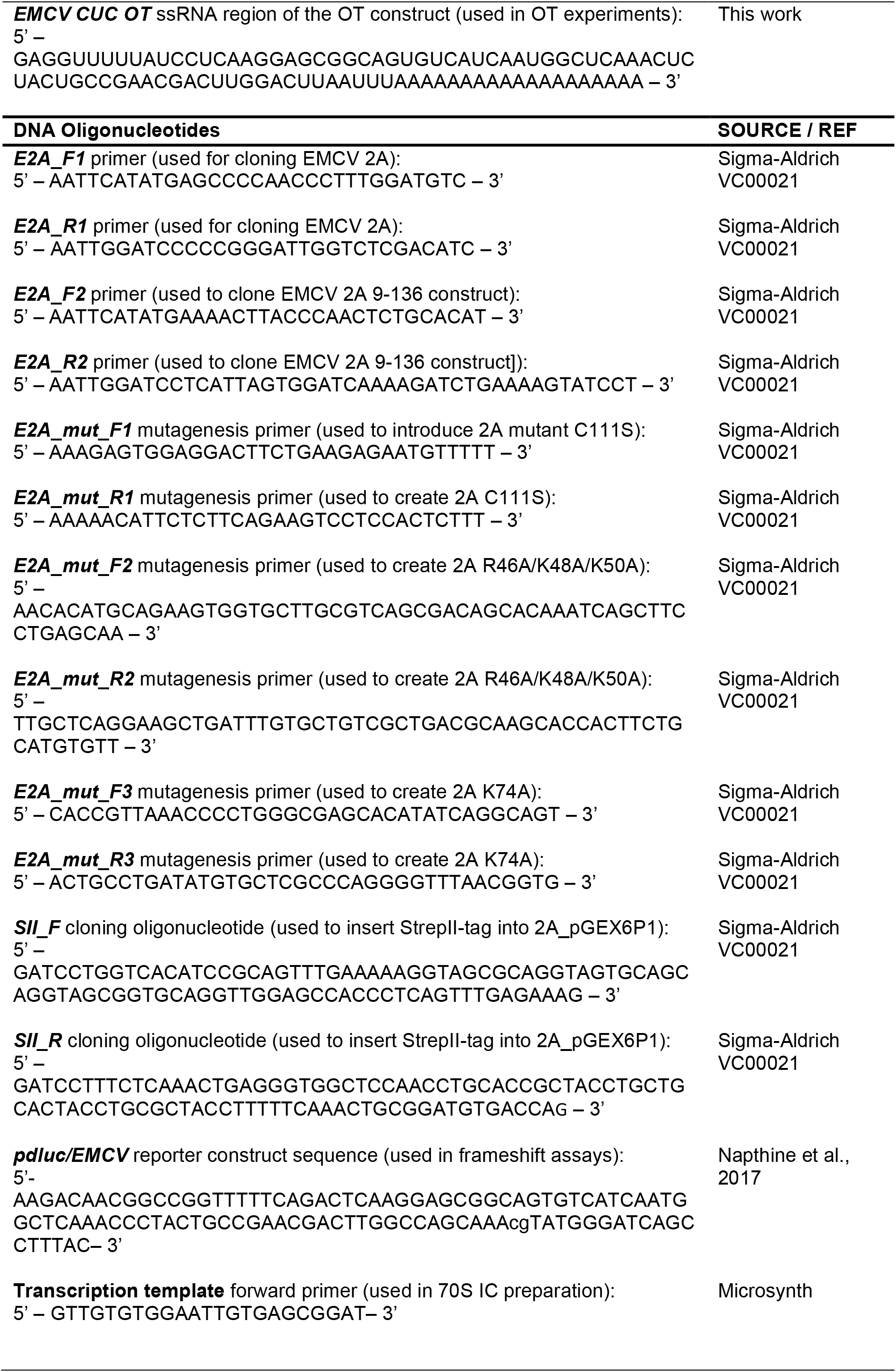
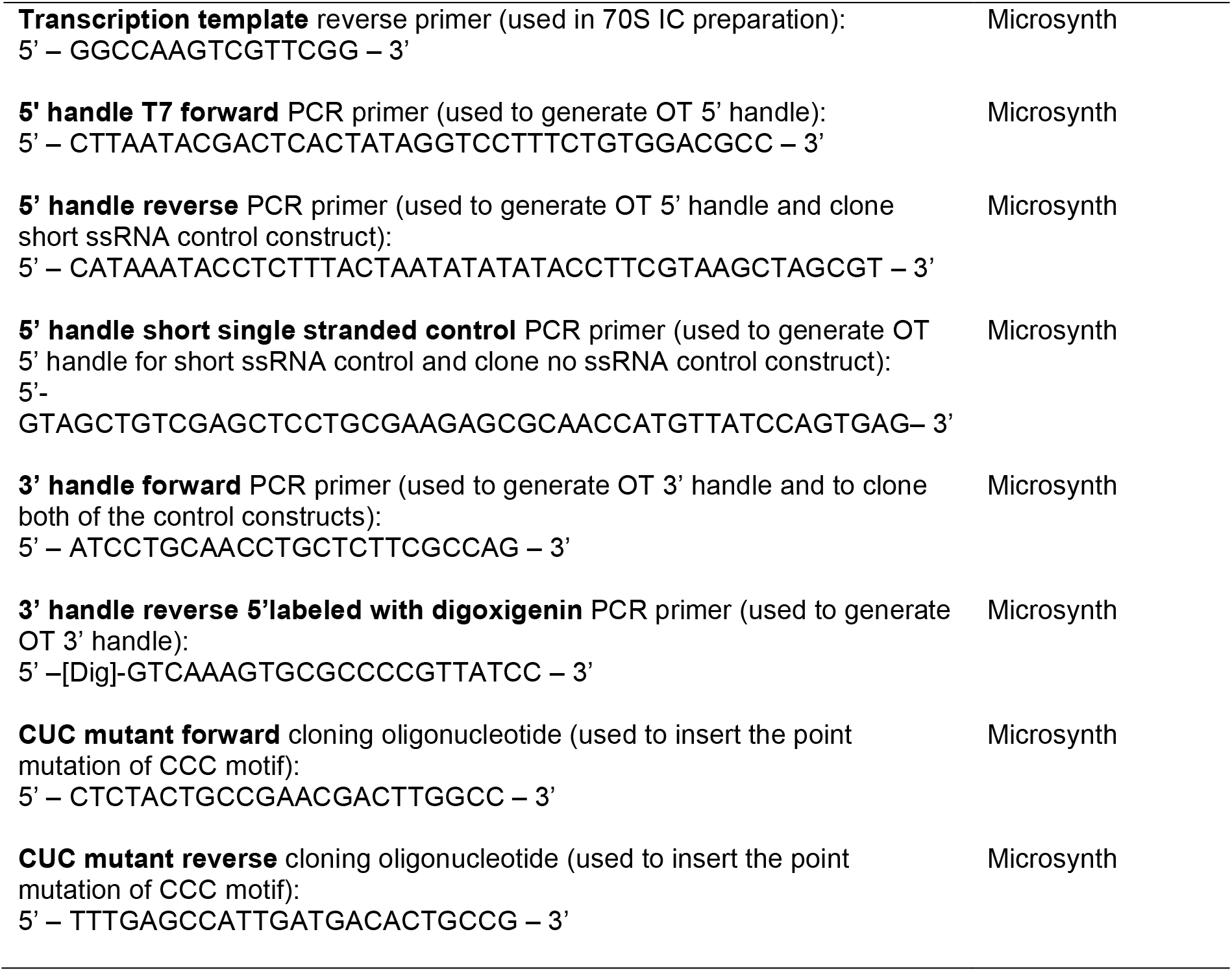
Oligonucleotide sequences. Related to **STAR Methods**.

#### Data and code availability

- Atomic coordinates and structure factors for the EMCV 2A X-ray crystal structure have been deposited in the wwPDB (XXXX).
- The 70S IC:2A cryo-EM map has been deposited in the EMDB (EMD-XXXX), and refined atomic coordinates accompanying this structure in the wwPDB (XXXX).
- Raw data (e.g. uncropped, unannotated gels, western blots, tables of force measurements, MST traces) corresponding to individual figure panels have been deposited in Mendeley Data (https://data.mendeley.com/XXXX).
- The force spectroscopy analysis scripts supporting the current study are available from Neva Caliskan on request.

### Experimental Model and Subject Details

All gene cloning, manipulation and plasmid propagation steps involving pGEX6P1 or pOPT vectors were carried out in *Escherichia coli* DH5α cells grown at 37 °C in 2 × TY or LB media supplemented with appropriate selection antibiotics as detailed in the text.

Recombinant proteins 2A, 2A_9-136_; C111S, 2A_R95A/R97A_, 2A_R46A/K48A/K50A_ and 2A_K73A_ were expressed in *E. coli* BL21 (DE3) pLysS cells grown in 2 × TY broth supplemented with 100 μg/mL ampicillin and 12.5 μg/mL chloramphenicol (37 °C, 200 rpm) until an OD_600nm_ of 0.6 – 1.0 was reached. Unless otherwise stated, expression was induced with 0.5 mM IPTG for either 4 h at 37 °C or overnight at 21 °C.

### Method Details

#### Cloning, protein expression and purification

EMCV 2A cDNA was amplified by PCR from previously described plasmid 2A_pGEX6P1 (Napthine et al., 2017) (primers E2A_F1 and E2A_R1; **Table 5**) and cloned into pOPTnH (Neidel et al., 2015) using NdeI and BamHI sites, thereby introducing a C-terminal GlySerLysHis_6_ tag. The 2A_9-136_ truncated construct was cloned in an identical way (primers E2A_F2 and E2A_R2; **Table 5**). The EMCV 2A R95A/R97A mutant was cloned into pOPTnH after PCR-amplification from a previously described 2A_pGEX6P1 construct containing these mutations (Napthine et al., 2017). Other EMCV 2A mutants were prepared by PCR mutagenesis, using either the wild-type EMCV 2A_pOPT or 2A_9-136__pOPT plasmids as templates, with the following primer pairs (C111S: E2A_mut_F1 and E2A_mut_R1; R46A/K48A/K50A: E2A_mut_F2 and E2A_mut_R2; K74A: E2A_mut_F3 and E2A_mut_R3; **Table 5**). To introduce an N-terminal StrepII-tag (_SII_-2A), annealed oligonucleotides encoding the StrepII-tag (SII_F and SII_R, Table 5) were inserted in-frame at the BamHI site of 2A_pGEX6P1. Proteins were expressed in *E. coli* BL21(DE3) pLysS cells. For selenomethionyl derivatisation (2ASeMet), protein was expressed in *E. coli* B834 cells, grown shaking (210 rpm, 37°C) in SeMet base media (Molecular Dimensions) supplemented with nutrient mix, 40 μg/mL L-selenomethionine and 100 μg/mL ampicillin. Expression was induced as above.

Cells were harvested by centrifugation (4,000 × g, 4°C, 20 min), washed once in ice-cold PBS and stored at −20°C. Pellets from four litres of culture were resuspended in cold lysis buffer (50 mM Tris-HCl pH 8.0, 500 mM NaCl, 30 mM imidazole, supplemented with 50 μg/mL DNase I and EDTA-free protease inhibitors) and lysed by passage through a cell disruptor at 24 kPSI (Constant Systems). Lysate was cleared by centrifugation (39,000 × g, 40 min, 4°C) prior to incubation (1 h, 4°C) with 4.0 mL of Ni-NTA agarose (Qiagen) pre-equilibrated in the same buffer. Beads were washed in batch four times with 200 mL buffer (as above, but without DNase or protease inhibitors) by centrifugation (600 × g, 10 min, 4°C) and re-suspension. Washed beads were pooled to a gravity column prior to elution over 10 column volumes (CV) with 50 mM Tris-HCl pH 8.0, 150 mM NaCl, 300 mM imidazole. Fractions containing 2A were pooled and dialysed (3K molecular weight cut-off (MWCO), 4°C, 16 h) against 1 L buffer A (50 mM Tris-HCl pH 8.0, 400 mM NaCl, 5.0 mM DTT) before heparin-affinity chromatography to remove contaminating nucleic acids. Samples were loaded on a 10 mL HiTrap Heparin column (GE Healthcare) at 2.0 mL/min, washed with two CV of buffer A and eluted with a 40% ⟶ 100% gradient of buffer B (50 mM Tris-HCl pH 8.0, 1.0 M NaCl, 5.0 mM DTT) over 10 CV. Fractions containing 2A were pooled and concentrated using an Amicon^®^ Ultra centrifugal filter unit (10K MWCO, 4,000 × g). Size exclusion chromatography was performed using a Superdex 75 16/600 column pre-equilibrated in 10 mM HEPES pH 7.9, 1.0 M NaCl, 5.0 mM DTT. Purity was judged by 4-20% gradient SDS-PAGE, and protein identity verified by mass spectrometry. Purified protein was used immediately or was concentrated as above (~ 7.0 mg/mL, 390 μM), snap-frozen in liquid nitrogen and stored at −80°C. Variants of 2A, including 2A_9-136;C111S_ and 2A_SeMet_ were purified identically to the wild-type protein. The StrepII-tagged variant (_SII_-2A) was expressed and purified using GST-affinity as previously described (Napthine et al., 2017). Following removal of the GST tag by 3C protease, _SII_-2A was further purified by Heparin affinity and size-exclusion chromatography as above.

#### Size-exclusion chromatography coupled to multi-angle light scattering (SEC-MALS)

Per experiment, 100 μL of protein was injected onto a Superdex 75 increase 10/300 GL column (GE Healthcare) pre-equilibrated with 20 mM Tris-HCl, 1.0 M NaCl (0.4 mL/min flow, 25°C). Experiments were performed with 5.2 mg/mL 2A (corresponding to a molar concentration of 290 μM). The static light scattering, differential refractive index, and the UV absorbance at 280 nm were measured in-line by DAWN 8+ (Wyatt Technology), Optilab T-rEX (Wyatt Technology), and Agilent 1260 UV (Agilent Technologies) detectors. The corresponding molar mass from each elution peak was calculated using ASTRA 6 software (Wyatt Technology).

#### Protein crystallization

Purified EMCV 2A was concentrated to 5.9 mg/ml in 10 mM HEPES pH 7.9, 1.0 M NaCl, 2.0 mM DTT. Diffraction-quality native 2A crystals were grown at 21°C by sitting-drop vapor diffusion against an 80 μL reservoir of 0.625 M (NH_4_)_2_SO_4_, 0.15 M tri-sodium citrate pH 5.7. Notably, crystal growth was only visible after 30 days. Drops were prepared by mixing 200 nL protein and 200 nL crystallization buffer. Selenomethionyl derivative 2A (2A_SeMet_) was concentrated to 5.7 mg/mL in 10 mM HEPES pH 7.9, 1.0 M NaCl, 2.0 mM DTT, and diffraction-quality 2A_SeMet_ crystals were grown as above against an 80 μL reservoir of 0.675 M (NH_4_)_2_SO_4_, 0.15 M tri-sodium citrate pH 5.7. Crystals were cryo-protected by the addition of 0.5 μL crystallization buffer supplemented with 20% v/v glycerol, prior to harvesting in nylon loops and flash-cooling by plunging into liquid nitrogen.

#### X-ray data collection, structure determination, refinement and analysis

Native datasets (**Table 1**) of 900 images were recorded at Diamond Light Source, beamline I03 (λ = 0.9796 Å) on a Pilatus 6M detector (Dectris), using 100% transmission, an oscillation range of 0.2° and an exposure time of 0.04 s per image. Data were collected at a temperature of 100 K. Data were processed with the XIA2 (Winter, 2009) automated pipeline, using XDS (Kabsch, 2010) for indexing and integration, and AIMLESS (Evans and Murshudov, 2013) for scaling and merging. Resolution cut-off was decided by a CC_1/2_ value ≥ 0.5 and an I/σ(I) ≥ 1.0 in the highest resolution shell (Karplus and Diederichs, 2012). For multiple-wavelength anomalous dispersion (MAD) phasing experiments, selenomethionyl derivative datasets were recorded at beamline I03 (peak λ = 0.9796 Å, 12656.0 eV; hrem λ = 0.9763, 12699.4 eV; inflexion λ = 0.9797 Å, 12655.0 eV). Data were processed as above using XIA2, XDS and AIMLESS. The structure was solved by three-wavelength anomalous dispersion analysis of the selenium derivative (space group *P*6_2_22) performed using the autoSHARP pipeline (Vonrhein et al., 2007), implementing SHELXD (Sheldrick, 2008) for substructure determination, SHARP for heavy-atom refinement and phasing, SOLOMON (Abrahams and Leslie, 1996) for density modification and ARP/wARP (Perrakis et al., 2001) for automated model building. This was successful in placing 503/573 (87%) residues in the asymmetric unit, which comprised four copies of the protein related by non-crystallographic symmetry (NCS). This initial model was then used to solve the native dataset by molecular replacement with Phaser (McCoy et al., 2007). The model was completed manually by iterative cycles of model-building using COOT (Emsley et al., 2010) and refinement with phenix.refine (Adams et al., 2010), using local NCS restraints and one TLS group per chain. Upon completion of model building, ISOLDE (Croll, 2018) was used to improve model geometry and resolve clashes prior to a final round of refinement using phenix.refine. MolProbity (Chen et al., 2010) was used throughout the process to evaluate model geometry. For the electrostatic potential calculations, partial charges were first assigned using PDB2PQR (Dolinsky et al., 2004), implementing PROPKA to estimate protein pKa values. Electrostatic surfaces were then calculated using APBS (Baker et al., 2001). Prior to designation of the “beta shell” as a new fold, structure-based database searches for proteins with similar folds to EMCV 2A were performed using PDBeFOLD (Krissinel and Henrick, 2004), DALI (Holm and Laakso, 2016) and CATHEDRAL (Redfern et al., 2007). Buried surface areas were calculated using PDBePISA (Krissinel and Henrick, 2007).

#### RNA folding prediction

The simRNAweb server (Magnus et al., 2016) was used for stem-loop and pseudoknot tertiary structure modelling of the EMCV stimulatory element. Experimentally-determined base-pairs were input as secondary structure restraints. Replica exchange Monte Carlo (REMC) simulated-annealing was performed with 10 replicas and 16000000 iterations per cycle. Trajectory files from eight independent simulations were concatenated and clustered, and all-atom PDB files was generated from the lowest energy state in each of the five most populous clusters. The 3D models presented **(Figure S3D)** represent the top cluster for pseudoknots and the top three clusters for stem-loops.

#### Electrophoretic Mobility Shift Assay (EMSA)

Synthetic RNA oligonucleotides (**Table 5**, IDT) were dissolved in distilled water. RNAs were labelled at the 5′ end with A647-maleimide or Cy5-maleimide conjugates (GE Healthcare) using the 5′ EndTag kit (Vector Labs) as directed by the manufacturer. For each binding experiment, a series of reactions were prepared on ice, each containing 1.0 μL 500 nM RNA, 1.0 μL serially diluted protein at concentrations of 320, 160, 80, 40, 20, 10, 5.0, and 2.5 μM in 10 mM HEPES pH 7.9, 1.0 M NaCl, 5.0 μL 2 × buffer (20 mM Tris-HCl pH 7.4, 80 mM NaCl, 4.0 mM magnesium acetate 2.0 mM DTT, 10% v/v glycerol, 0.02% w/v bromophenol blue, 200 μg/mL porcine liver tRNA, 800 U /mL SUPERase-In [Invitrogen]) and 3.0 μL distilled water. This gave final binding reactions of 10 μL with 50 nM RNA, 1 × buffer, a salt concentration of ~ 140 mM and proteins at concentrations of 32, 16, 8.0, 4.0, 2.0, 1.0, 0.5 and 0.25 μM. Samples were incubated at 37°C for 20 min prior to analysis by native 10% acrylamide/TBE PAGE (25 min, 200 V constant). Gels were scanned with a Typhoon FLA-7000 (GE) using the 635 nm laser / R670 filter.

#### Isothermal Titration Calorimetry (ITC)

ITC experiments were performed at 25°C using an automated MicroCal PEAQ-ITC platform (Malvern Panalytical). Proteins and synthetic RNA oligonucleotides (IDT) were dialysed extensively (24 h, 4°C) into buffer (50 mM Tris-HCl pH 7.4, 400 mM NaCl) prior to experiments. RNA (52 μM) was titrated into protein (5 μM) with 1 × 0.4 μL injection followed by 12 × 3.0 μL injections. Control titrations of RNA into buffer, buffer into protein and buffer into buffer were also performed. Data were analysed using the MicroCal PEAQ-ITC analysis software (Malvern Panalytical) and fitted using a one-site binding model.

#### Microscale Thermophoresis (MST)

For RNA-binding experiments, synthetic EMCV RNA variants **(Table 5)** were dissolved in distilled water and labelled at the 5’ end with Dylight 650 maleimide conjugates (Thermo Scientific) using the 5′ EndTag kit (Vector Labs) as directed by the manufacturer. For each binding experiment, RNA was diluted to 10 nM in MST buffer (50 mM Tris-HCl pH 7.8, 150 mM NaCl, 10 mM MgCl_2_, 2 mM DTT supplemented with 0.05% Tween 20) and a series of 16 tubes with 2A dilutions were prepared on ice in MST buffer, producing 2A ligand concentrations ranging from 0.00015 to 5 μM for EMCV RNA 2-6 and 0.00006 to 20 μM for EMCV RNA1. For the measurement, each ligand dilution was mixed with one volume of labelled RNA, which led to a final concentration of 5.0 nM labelled RNA. The reaction was mixed by pipetting, incubated for 10 min followed by centrifugation at 10,000 × g for 10 min. Capillary forces were used to load the samples into Monolith NT.115 Premium Capillaries (NanoTemper Technologies). Measurements were performed using a Monolith NT.115Pico instrument (NanoTemper Technologies) at an ambient temperature of 25°C. Instrument parameters were adjusted to 5% LED power, medium MST power and MST on-time of 10 seconds. An initial fluorescence scan was performed across the capillaries to determine the sample quality and afterwards 16 subsequent thermophoresis measurements were performed. To determine binding affinities, data of at least two independently pipetted measurements were analysed for the fraction bound (MO.Affinity Analysis software, NanoTemper Technologies). For the non-binders, since the maximum amplitude would numerically be zero, deltaFnorm values were divided by the average maximum amplitude of the dataset to plot fraction bound. Data were fitted to the Kd model using MO.Affinity Analysis software (NanoTemper) and were plotted using Prism 8.0.2 (GraphPad).

Conjugation of a fluorescent label to the surface-exposed cysteine residue (C111) observed in the 2A crystal structure **(Figure 1E)** provided a convenient way of studying binding to multiple unlabelled targets by MST, in such a way that the observed affinities would be directly comparable. For this experiment, EMCV 2A protein was labelled using the Protein Labelling Kit RED-Maleimide (NanoTemper Technologies) according to the manufacturer’s instructions. In brief, 2A protein was diluted in a buffer containing 10 mM HEPES pH 7.9, 1.0 M NaCl and dye was mixed at a 1:3 molar ratio at room temperature for 30 min in the dark. Unreacted dye was removed on a spin gel filtration column equilibrated with 10 mM HEPES pH 7.9, 1.0 M NaCl. The labelled 2A protein was diluted to 10 nM in MST buffer. Synthetic EMCV RNA variants were used in dilutions ranging from 0.0008 to 26 μM for RNA 1 and 0.00003 to 1 μM for RNA 2-6. For the measurement, each RNA ligand dilution was mixed with one volume of labelled protein 2A, which led to a final concentration of protein 2A of 5.0 nM. Similar experiments were conducted with ribosomes in MST buffer, with ligand concentrations ranging between 0.00002 to 0.4 μM for 40S and 60S, 0.00003 to 1 μM for 30S, 0.000027 to 0.9 μM for 50S, 0.0008 to 1.375 μM for empty 70S and 0.000003 to 0.1 μM for 70S IC. The measurements were performed as described above.

#### Preparation of constructs for optical tweezer experiments

DNA encoding the frameshifting sequence of EMCV was inserted into plasmid pMZ_lambda_OT using PCR and subsequent Gibson assembly. This plasmid contains the ColE1 origin, ampicillin resistance, ribosome binding site and two 2 kbp handle regions derived from lambda phage DNA (5′ and 3′ handle). For the generation of the mutant plasmid, PCR and blunt-end ligation was used to mutate the CCC triplet in the EMCV stem-loop to CUC. Control constructs (see below) were prepared the same way as mutant constructs. For the control construct without any single-stranded RNA region, a PCR reaction using the EMCV wild-type (CCC) construct as template was conducted with 3’ handle forward oligonucleotide and 5’ handle reverse oligonucleotide as primers **(Table 5)**. For the control construct with a short single-stranded RNA region, a PCR reaction using the EMCV wild-type (CCC) construct as a template was conducted with the 3’ handle forward oligonucleotide and the 5’ handle RBS reverse oligonucleotide as primers **(Table 5)**. After the PCR, the linear products were blunt-end ligated to yield the control constructs. Wild-type and mutant plasmids were subsequently used to generate construct suitable for optical tweezer measurements consisting of the EMCV frameshifting sequence flanked by the 2 kbp long handle regions. Three pairs of primers for PCR were designed allowing the amplification of the *in vitro* transcription template and 5′ and 3′ handles. Subsequently, PCR reactions generated 5′ and 3′ handles and a long template for *in vitro* transcription. The 3′ handle was labelled during PCR using a 5′ digoxigenin-labelled reverse primer. The 5′ handle was labelled with Biotin-16-dUTP at the 3′ end following PCR using T4 DNA polymerase. RNA was transcribed from templates for *in vitro* transcription using T7 RNA polymerase. RNA and both DNA handles (5′ and 3′) were annealed together in a mass ratio 1:1:1 (5 μg each) by incubation at 95 °C for 10 min, 62 °C for 1 hour, 52 °C for 1 hour and slow cooling to 4 °C in a buffer containing 80% formamide, 400 mM NaCl, 40 mM HEPES, pH 7.5, and 1 mM EDTA (Stephenson et al., 2014). Following annealing, the samples were concentrated by ethanol precipitation, the pellets resuspended in 40 μL RNase-free water, split into 4 μL aliquots and stored at –20 °C.

#### Optical tweezers data collection and analysis

Optical tweezers experiments were performed using a commercial dual-trap platform equipped with a microfluidics system (C-trap, Lumicks). For the experiments, optical tweezers (OT) constructs from above were mixed with 1 μL of polystyrene beads coated with antibodies against digoxigenin (0.1% v/v suspension, Ø 1.76 μm, Lumicks), 5 μL of measurement buffer (20 mM HEPES, pH 7.6, 300 mM KCl, 5 mM MgCl_2_, 5 mM DTT and 0.05% Tween) and 0.5 μL of RNase inhibitors. The mixture was incubated for 20 min at room temperature in a final volume of 10.5 μL, and subsequently diluted by addition of 0.5 mL measurement buffer. Separately, 0.8 μL of streptavidin-coated polystyrene beads (1% v/v suspension, Ø 2 μm, Lumicks) was supplemented with 1 mL of measurement buffer, the flow cell was washed with the measurement buffer and suspensions of both streptavidin beads as well as the complex of OT construct with anti-digoxigenin beads were introduced into the flow cell. Per experiment, an anti-digoxigenin (AD) bead and a streptavidin (SA) bead were optically trapped and brought into close proximity to allow the formation of a tether in between. The beads were moved apart (unfolding) and back together (refolding) at constant speed (0.05 μm/s) to yield the force-distance (FD) curves. The stiffness was maintained at 0.31 and 0.24 pN/nm for trap 1 (AD bead) and trap 2 (SA bead), respectively. For experiments with 2A protein experiments, protein was diluted to 300 nM in measurement buffer and added to the buffer channel of the optical tweezer flow cell. FD data was recorded at a rate of 78000 Hz. To ensure that the observed effects were indeed a result of interaction with the studied RNA region and not a non-specific binding to handle regions, we also employed constructs containing either no single-stranded RNA sequence (No ssRNA control) or a short (43 nt) stretch of single-stranded RNA (short ssRNA) (https://data.mendeley.com/XXXX).

Afterwards, the data were down sampled by a factor of 20 and filtered with a Butterworth filter (0.05 filtering frequency, filter order 2) using Matlab. Individual unfolding/refolding steps were manually marked by custom written Matlab scripts and the unfolding force and step length were calculated based on the marked coordinates. Unfolding and refolding steps were then plotted for each sample as heat maps. After that, individual traces were manually sorted into “state” groups based on similar trace and steps character. FD curves were fitted using a custom written Python script, which is based on Pylake package provided by Lumicks (https://lumicks-pylake.readthedocs.io/). The fitting procedure was done as described in (Mukhortava et al., 2019). In brief, first, a fully folded part (until the first detectable unfolding step) was fitted with a worm-like chain model (WLC) (Odijk, 1995; Wang et al., 1997) to determine the persistence length (dsL_P_) of the tether while the contour length (dsL_C_) parameter was held fixed at 1256 nm (± 1%; 4110 bp*0.305 nm/bp and 4 ss*0.59 nm/ss) (Zhang et al., 2019). The (partially) unfolded parts of FD curve were then fitted by a model comprising of WLC (describing the folded double stranded handles) and freely jointed chain (FJC) model (Smith et al., 1996; Wang et al., 1997) (describing the unfolded single stranded parts). For fitting of the unfolded regions, parameters extracted from the fully folded part fitting (dsL_P_, dsL_C_, dsK) were used and fixed in WLC part of the combined model. Persistence length of the single stranded part (ssLP) was fixed at 1 nm while contour length (ssLC) of the single stranded part together with the single stranded stretch modulus (ssK) were optimized. The FD curves were plotted using Prism 8.0.2 (GraphPad), for final plots the data were filtered with Butterworth filter (0.005 filtering frequency, filter order 4). The RNAstructure software (version 6.2) was used for the prediction of the EMCV RNA element secondary structure (Bellaousov et al., 2013).

#### Eukaryotic ribosomal subunit purification

40S and 60S subunits were purified from untreated rabbit reticulocyte lysate (Green Hectares) as previously described^44^. Briefly, ribosomes were pelleted by centrifugation (4°C, 270,000 × g, 4.5 h) and resuspended in 20 mM Tris-HCl pH 7.5, 4.0 mM MgCl_2_, 50 mM KCl, 2.0 mM DTT. Following treatment with 1.0 mM puromycin and addition of KCl to 0.5 M, 40S and 60S subunits were separated by centrifugation (4°C, 87,000 × g, 16 h) through a sucrose density gradient (10 ⟶ 30% sucrose in 20 mM Tris-HCl pH 7.5, 2.0 mM DTT, 4.0 mM MgCl_2_, 0.5 M KCl). After analysis by SDS-PAGE, uncontaminated fractions were pooled, and exchanged into 20 mM Tris-HCl pH 7.5, 100 mM KCl, 2.0 mM MgCl_2_, 2.0 mM DTT, 250 mM sucrose using Amicon centrifugal concentrators (4°C,100K MWCO). Ribosome subunits were snap-frozen in liquid nitrogen and stored at −80°C until required.

#### Ribosome binding assays

Assays were conducted in 50 mM Tris-acetate pH 7.5, 150 mM potassium acetate, 5.0 mM magnesium acetate, 0.25 mM spermidine, 10 mM DTT, 0.1 % v/v Triton X-100. Per 60 μL binding reaction, ribosome subunits were diluted to a final concentration of 0.4 μM, and 2A protein was added in excess to a final concentration of 2.4 μM. Twenty microlitres of this mixture was retained for SDS-PAGE analysis of the ‘input’. The remaining 40 μL was incubated at room temperature for 20 min prior to application to a S200-HR size-exclusion microspin column (Cytiva) that had been pre-equilibrated (4 × 500 μL) in the above buffer by resuspension and centrifugation (300 × g, 30 s). Immediately after application, the eluate was collected by centrifugation (300 × g, 60 s).

#### Western blot

Samples were analysed by 4–20% gradient SDS-PAGE and transferred to a 0.2 μm nitrocellulose membrane. All subsequent steps were carried out at room temperature. Membranes were blocked (5% w/v milk, PBS, 1 h) before incubation (1 h) with primary antibodies in 5% w/v milk, PBS, 0.1% v/v Tween-20. Membranes were washed three times with PBS, 0.1% v/v Tween-20 prior to incubation (1 h) with IRDye fluorescent antibodies in 5% w/v milk, PBS, 0.1% v/v Tween-20. After three washes in PBS, 0.1% v/v Tween-20 and a final rinse in PBS, membranes were imaged using an Odyssey CLx Imaging System (LI-COR). Antibodies used were rabbit polyclonal anti-2A (Napthine et al., 2017) (1/1000); mouse monoclonal anti-RPS6 (1/1000, clone A16009C, BioLegend); mouse monoclonal anti-RPL4 (1/1000, clone 4A3, Sigma); goat anti-rabbit IRDye 800 CW (1/10,000, LI-COR) and goat anti-mouse IRDye 680LT (1/10,000, LI-COR).

#### *In vitro* transcription

For *in vitro* frameshifting assays, we cloned a 105 nt DNA fragment (pdluc/EMCV, **Table 5**) containing the EMCV slippery sequence flanked by 12 nt upstream and 86 nt downstream into the dual luciferase plasmid pDluc at the XhoI and BglII sites (Fixsen and Howard, 2010). This sequence was inserted between the Renilla and firefly luciferase genes such that firefly luciferase expression is dependent on −1 PRF. Wild-type or mutated frameshift reporter plasmids were linearized with FspI and capped run-off transcripts generated using T7 RNA polymerase as described (Powell et al., 2008). Messenger RNAs were recovered by phenol/chloroform extraction (1:1 v/v), desalted by centrifugation through a NucAway Spin Column (Ambion) and concentrated by ethanol precipitation. The mRNA was resuspended in water, checked for integrity by agarose gel electrophoresis, and quantified by spectrophotometry.

Messenger RNAs for 70S IC preparation (EMCV_IC, **Table 5**) were produced from a 117 nt long DNA fragment containing the EMCV frameshift site flanked by the bacterial 5′ UTR with Shine-Dalgarno sequence and 18 nt downstream region of the putative structure.

5′GGGAAUUCAAAAAUUGUUAAGAAUUAAGGAGAUAUACAU<u>AUG</u>GA**GGUUUUU**AUCACUCAA GGAGCGGCAGUGUCAUCAAUGGCUCAAACCCUACUGCCGAACGACUUGGCCAGATCT 3′(slippery sequence in bold, initiation codon underlined)

This sequence was PCR amplified and *in vitro* transcribed using T7 RNA polymerase (produced in-house). Messenger RNAs were purified using the Qiagen RNeasy midiprep kit according to the manufacturer’s protocols. The mRNAs were eluted in RNAse-free water, integrity and purity was checked by gel electrophoresis and quantified by spectrophotometry.

#### 70S initiation complex preparation

Ribosomes, translation factors, and tRNAs were of *E. coli* origin. Total *E. coli* tRNA was from Roche, and oligonucleotides were from Microsynth. 70S ribosomes from MRE600, EF-Tu, EF-G, IF1, IF2 and IF3 were purified from *E. coli* (Milon et al., 2007). fMet-tRNAfMet was prepared and aminoacylated according to published protocols (Kothe et al., 2006; Rodnina et al., 1994). Aminoacylated fMet-tRNAfMet was purified by reversed-phase HPLC on a Wide Pore C5 (10

μM particle size 10 mm × 25 cm) column (Sigma Aldrich). To prepare initiation complexes, 70S ribosomes (1 μM) were incubated with a three-fold excess of an EMCV model mRNA (EMCV_IC, **Table 5**) encoding for 5′…AUGGA**GGUUUUU**AUC…3′ (slippery sequence in bold) and a 1.5-fold excess each of IF1, IF2, IF3, and fMet-tRNA^fMet^ in buffer A (50 mM Tris-HCl pH 7.5, 70 mM NH4Cl, 30 mM KCl, 7 mM MgCl_2_) supplemented with GTP (1 mM) for 30 min at 37°C. 70S initiation complexes were purified by centrifugation through a 1.1 M sucrose cushion in buffer A. Before grid preparation, initiation complexes were additionally purified on Sephacryl S-300 gel filtration microspin columns.

#### Frameshifting assays (*In vitro* translation)

Messenger RNAs were translated in nuclease-treated rabbit reticulocyte lysate (RRL) or wheat germ (WG) extracts (Promega). Typical reactions were composed of 90% v/v RRL, 20 μM amino acids (lacking methionine) and 0.2 MBq [^35^S]-methionine and programmed with ∼50 μg/mL template mRNA. Reactions were incubated for 1 h at 30°C. Samples were mixed with 10 volumes of 2× Laemmli’s sample buffer, boiled for 3 min and resolved by SDS-PAGE. Dried gels were exposed to a Storage Phosphor Screen (PerkinElmer) and the screen scanned in a Typhoon FLA7000 using phosphor autoradiography mode. Bands were quantified using ImageQuant™TL software. The calculations of frameshifting efficiency (%FS) took into account the differential methionine content of the various products and %FS was calculated as % −1FS = 100 × (IFS/MetFS) / (IS/MetS + IFS/MetFS). In the formula, the number of methionines in the stop and frameshift products are denoted by MetS, MetFS respectively; while the densitometry values for the same products are denoted by IS and IFS respectively. All frameshift assays were carried out a minimum of three times.

Ribosomal frameshift assays in *E. coli* employed a coupled T7/S30 *in vitro* translation system (Promega). A ~450 bp fragment containing the EMCV PRF signal (or mutant derivative) was prepared by PCR from plasmid pDluc/EMCV (Napthine et al., 2017) and cloned into the BamHI site of the T7-based, *E. coli* expression vector pET3xc (Studier et al., 1990). T7/S30 reaction mixes were prepared according to the manufacturer’s instructions (50 μL volumes), including 10 μCi ^35^S methionine, supplemented with plasmid DNA (4 μg) and incubated at 37 °C for 90 mins. Reactions were precipitated by addition of an equal volume of acetone, dissolved in Laemmli’s sample buffer and aliquots analysed by SDS-PAGE. PRF efficiencies were calculated as above.

#### Cryo-EM specimen preparation

Initiated 70S ribosomes in 50 mM Tris-HCl pH 7.5, 70 mM NH4Cl, 30 mM KCl, 7 mM MgCl_2_ were diluted tenfold into 20 mM HEPES pH 7.5, 100 mM potassium acetate, 1.5 mM MgCl_2_, 2.0 mM DTT. 2A protein was dialysed (3K MWCO, 4°C, 16 h) into the same buffer. Crosslinking reactions of 50 μL comprising 75 nM ribosomes, 3.0 μM 2A and 2.0 mM bis(sulfosuccinimidyl)suberate (BS3) were performed on ice (30 min) immediately prior to grid preparation. Quantifoil R 2/2 400-mesh copper supports were coated with an additional ~ 60 Å layer of amorphous, evaporated carbon by flotation (Passmore and Russo, 2016), and thoroughly dried before use. Grids were made hydrophilic by glow-discharge in air for 30 s. Three microliters of crosslinking reaction was applied to grids which were then blotted for 4.5 s and vitrified by plunging into liquid ethane using a Vitrobot MK IV (FEI) at 4°C, 100% relative humidity.

#### Cryo-EM data collection and processing

Micrographs were collected at the BiocEM facility (Department of Biochemistry, University of Cambridge) on a Titan Krios microscope (FEI) operating at 300 kV and equipped with a Falcon III detector **(Table 4)**. At 75,000 × magnification, the calibrated pixel size was 1.07 Å / pixel. Per 0.6 s acquisition in integration mode, a total exposure of 54.4 e^−^/ Å^2^ was fractionated over 23 frames with applied defocus of –1.5, –1.8, –2.1, –2.4, –2.7 and –3.0 μm. EPU software was used for automated acquisition with five images per hole. After manual inspection, 5730 micrographs were used in subsequent image processing.

Movie frames were aligned and a dose-weighted average calculated with MotionCor2 (Zheng et al., 2017) The contrast transfer function (CTF) was estimated using CtfFind4 (Rohou and Grigorieff, 2015). All subsequent image-processing steps were carried out in RELION 3.1 (Zivanov et al., 2018) **(Fig S6)** and all reported estimates of resolution are based on the gold standard Fourier shell correlation (FSC) at 0.143, and the calculated FSC is derived from comparisons between reconstructions from two independently refined half-sets. Reference-free autopicking of 820,475 particles was performed using the Laplacian-of-Gaussian function (200 - 250 Å diameter). Particles were initially downscaled threefold and extracted in a 150-pixel box. Two rounds of 2D classification (into 100 and 200 classes, respectively) were used to clean the dataset to 750,029 ‘good’ particles. An initial model was generated from a PDB file of a 70S elongation-competent ribosome (PDB ID 5MDZ) and low-pass filtered to 80 Å resolution. The initial 3D refinement (6.5 Å resolution) showed clear evidence for at least one copy of 2A adjacent to the factor binding site on the 30S subunit. At this stage, two rounds of focussed classification with signal subtraction were performed (6 classes) to separate particles based on additional density near i) the factor binding site and ii) the mRNA entry channel/helicase. The former was successful and 289,741 particles containing three copies of 2A were rescaled to full size and extracted in a 450-pixel box. Following initial 3D refinement, creation of a 15 Å low-pass filtered mask (five-pixel extension and five-pixel soft edge) and post-processing, a reconstruction of 2.93 Å was achieved. After per-particle CTF refinement and polishing, this was increased to 2.50 Å. With the increased angular accuracy provided by the fully rescaled data, focussed classification with signal subtraction and local angular searches was performed again to separate particles based on 2A occupancy at the factor binding site. The final reconstruction (2.66 Å) from 120,749 particles revealed three copies of 2A bound with full occupancy. Calculation of a local resolution map revealed additional low-resolution density adjacent to the beak of the 30S head. Subsequent focussed classification with signal subtraction and refinement confirmed that this was a fourth copy of 2A bound, present in 73,059 particles.

To build the model, the atomic coordinates for a 70S initiation complex (5MDZ) and three copies of chain A from the 2A crystal structure (above) were docked as rigid bodies into the EM map. Local rebuilding was performed iteratively in COOT (Emsley et al., 2010) and the models refined using phenix real-space refine (Adams et al., 2010) implementing reference model restraints to preserve geometry.

#### Visualisation of structural data

All structural figures depicting crystallographic data (cartoon, stick and surface representations) were rendered in PyMOL (Schrödinger LLC). Structural figures of EM maps with docked components were rendered in ChimeraX (Pettersen et al., 2004).

### Quantification and Statistical Analysis

#### Crystallographic and cryo-EM data

Crystallographic calculations (e.g. integration, scaling, merging) were performed as described in methods text, using the default software parameters unless otherwise stated. Processing and refinement statistics are detailed in **Table 1**. EM image processing (e.g. motion correction, CTF estimation, particle picking, 2D/3D classification, refinement, CTF-refinement and particle polishing) was performed as described in methods text, using default software parameters unless otherwise stated. Processing and refinement statistics are detailed in **Table 4**.

#### Force spectroscopy data

Force spectroscopy data processing and analysis was performed as described in STAR Methods text, using custom written Matlab and Python scripts, which were based on Pylake package provided by Lumicks.

#### Binding affinity data (ITC and MST)

Presented binding isotherms were analysed using the MicroCal PEAQ-ITC analysis software (Malvern Panalytical) and fitted using a one-site binding model. Presented traces were representative of two independent titrations. MST data presented are fraction bound values. Data were fitted to the Kd model using MO.Affinity Analysis software (NanoTemper) and were plotted using Prism 8.0.2 (GraphPad). Presented traces were representative of at least two independent titrations.

## Author contributions

C.H.H. and S.N. cloned expressed and purified proteins and performed all biochemical experiments. C.H.H. and S.C.G performed crystallography experiments. A.K. and N.C. performed MST experiments and analyses. L.P. performed single-molecule experiments and analyses. C.H.H. prepared cryo-EM grids, and collected and processed cryo-EM data. C.H.H., S.N., N.C. and I.B. wrote the manuscript with contributions from all authors.

## Acknowledgements

We thank Dima Chirgadze, Steve Hardwick and Lee Cooper at the BiocEM facility for assistance with CryoEM data acquisition. We thank Ann Mukhortova and Bärbel Lorenz (Lumicks AG) for expert technical assistance in optical tweezer data collection. We thank Matthias Zimmer for his participation in the initial phase of the work. We thank Prof. Marina V. Rodnina for providing expression constructs for bacterial 70S translation complexes. We thank Trevor Sweeney for providing ribosomes from rabbit reticulocyte lysate, and assistance with 40S and 60S purification. A Titan V graphics card used for this research was donated by the NVIDIA Corporation. Remote synchrotron access was supported in part by the EU FP7 infrastructure grant BIOSTRUCT-X (Contract No. 283570). We thank the staff of Diamond Light Source beamline I03 for assistance with crystal screening and data collection. We thank Janet Deane for assistance with SEC-MALS experiments. Part of this work was carried out in the laboratory of V. Ramakrishnan, who was funded by the UK Medical Research Council (MC_U105184332), and a Wellcome Trust Senior Investigator award (WT096570). We are grateful to Vish Chandrasekaran, Jailson Brito Querido, Sebastian Kraatz and Chris Rae for helpful discussions. We thank Tatyana Koch for expert technical assistance in ribosome purifications and translation initiation experiments. CHH and SN are funded by a Wellcome Trust Investigator Award (202797/Z/16/Z) to IB. AEF is supported by Wellcome Trust (106207/Z/14/Z) and European Research Council (646891) grants to AEF. SCG is funded by a Sir Henry Dale fellowship (098406/Z/12/B) funded by the Wellcome Trust and the Royal Society. NC, LP and AK are supported by the Helmholtz Association.

## Key Resources Table

*NB. “SII” denotes a StrepII tag and “6H” denotes a His6 tag. These descriptors are positioned before or after a gene/protein name based on whether tag is N- or C-terminal*

| REAGENT or RESOURCE | SOURCE | IDENTIFIER |
| --- | --- | --- |
| **Antibodies** |  |  |
| rabbit polyclonal anti-2A | Napthine et al., 2017 | N/A |
| mouse monoclonal anti-RPS6, clone A16009C | BioLegend | 691802 |
| mouse monoclonal anti-RPL4, clone 4A3 | Sigma-Aldrich | WH0006124M1 |
| goat anti-rabbit IRDye 800 CW | LI-COR | 926-32211 |
| goat anti-mouse IRDye 680LT | LI-COR | 926-68020 |
| **Bacterial and Virus Strains** |  |  |
| *E. coli* DH5α | Thermo Fisher Scientific | 18258012 |
| *E. coli* BL21 star pLysS | Thermo Fisher Scientific | C602003 |
| **Chemicals, Peptides, and Recombinant Proteins** |  |  |
| Protease Inhibitor Cocktail (EDTA-free) | Sigma-Aldrich | 11836170001 |
| Trizma hydrochloride | Sigma-Aldrich | T3253 |
| Potassium chloride | Merck | 529552-1KG |
| Magnesium chloride | Carl Roth | 2189.1 |
| HEPES | Carl Roth | HN78.3 |
| Sodium chloride | Sigma-Aldrich | 31424 |
| Glycerol | Sigma-Aldrich | G5516 |
| Sucrose | Fisher | S/8600/60 |
| Imidazole | Sigma-Aldrich | I5513 |
| Formamide | Sigma-Aldrich | 11814320001 |
| TEMED | Sigma-Aldrich | T9281 |
| Ammonium persulfate (APS) | Sigma-Aldrich | A3678 |
| ProtoGel (30% Solution at 37.5:1 acrylamide/methylene bisacrylamide Ratio) | Geneflow | A2-0072 |
| Urea | VWR chemicals | 28877.260 |
| KOD Hot Start DNA Polymerase | Merck | 71086 |
| Ethidium Bromide | Sigma-Aldrich | E1510 |
| Ni-NTA Agarose | QIAGEN | 30210 |
| bis(sulfosuccinimidyl)suberate (BS3) cross-linker | Thermo Fisher Scientific | 21580 |
| Instant Blue (coomassie) protein gel stain | Expedeon | ISB1L |
| Imperial (coomassie) protein gel stain | Thermo Scientific | 24615 |
| Dithiothreitol (DTT) | Melford Labs | D11000 |
| DNAse I from bovine pancreas | Sigma-Aldrich | D4527 |
| Deoxynucleoside Triphosphate Set | Roche | 11969064001 |
| Ribonucleoside Triphosphate Set | Roche | 11277057001 |
| m7G(5')ppp(5')G RNA Cap Structure Analog | NEB Biolabs | S1404S |
| T7 RNA Polymerase | Invitrogen | 18033100 |
| SP6 RNA Polymerase | Roche | 10810274001 |
| RNaseOUT™ Recombinant Ribonuclease Inhibitor | Invitrogen | 100000840 |
| Rabbit Reticulocyte Lysate, Nuclease-Treated | Promega | L4960 |
| EasyTag™ L-[35S]-Methionine, 5mCi (185MBq), Stabilized Aqueous Solution | PerkinElmer | NEG709A005MC |
| [Methyl-14C] Methylated, Protein Molecular Weight Markers | PerkinElmer | NEC811005UC |
| TWEEN 20 BioXtra | Sigma-Aldrich | P7949 |
| T4 DNA Polymerase | NEB | M0203S |
| Biotin-16-dUTP | Roche | 11093070910 |
| RNase Inhibitor | Molox | 1000379515 |
| Pyrophosphatase, thermostabile, inorganic | NEB | M0296L |
| PrimeSTAR GXL DNA Polymerase | Takara Bio Clontech | R050A |
| DyLight™ 680 Maleimide | ThermoFisher Scientific | 46618 |
| Protein Labelling Kit RED-Maleimide | Nanotemper | MO-L014 |
| Cy5 Maleimide mono-reactive dye | Cytiva | PA15131 |
| 5' EndTag labelling kit | Vector Labs | MB-9001 |
| Recombinant protein: 2A<sub>-6H</sub> | This work | N/A |
| Recombinant protein: SII-2A | This work | N/A |
| Recombinant protein: 2A<sub>-6H</sub> 9-136; C111S | This work | N/A |
| Recombinant protein: 2A<sub>-6H</sub> 9-136; C111S | This work | N/A |
| Recombinant protein: 2A<sub>-6H</sub> R95A/R97A | This work | N/A |
| Recombinant protein: 2A<sub>-6H</sub> R46A/K48A/K50A | This work | N/A |
| Recombinant protein: 2A<sub>-6H</sub> K73A | This work | N/A |
| **Critical Commercial Assays** |  |  |
| N/A |  |  |
| **Deposited Data** |  |  |
| Mendeley raw data (e.g. uncropped gels, blots, raw MST and force spectroscopy traces) | This work | XXXX |
| EMCV 2A crystal structure | This work | PDB: XXXX |
| 70S-mRNA-tRNA<sup>fMet</sup>-2A complex cryo-EM structure | This work | PDB: XXXX |
| 70S-mRNA-tRNA<sup>fMet</sup>-2A complex cryo-EM map | This work | EMD: XXXX |
| 70S-mRNA-tRNA<sup>fMet</sup> complex (initiation complex), starting model for refinement | James et al., 2016 | PDB: 5MDZ |
| 70S pre-translocation complex with EF-G (used for Figure 6) | Brilot et al., 2013 | PDB: 4V7D |
| 70S complex with EF-Tu (used for Figure 6) | Fislage et al., 2018 | PDB: 5WE6 |
| 70S complex with dityromycin (used for Figure S7) | Bulkley et al., 2014 | PDB: 4WQU |
| **Experimental Models: Cell Lines** |  |  |
| N/A |  |  |
| **Experimental Models: Organisms/Strains** |  |  |
| N/A |  |  |
| **Oligonucleotides** |  |  |
| RNA and DNA sequences, with details of end-labelling | This work | See **Table 5** |
| **Recombinant DNA** |  |  |
| 2A_pOPTnH (bacterial expression). Used to make protein:<br>2A<sub>-6H</sub> (wild-type) | This work | N/A |
| 2A_pOPTnH R95A/R97A (bacterial expression). Used to make protein:<br>2A<sub>-6H</sub> R95A/R97A | This work | N/A |
| 2A_pOPTnH R46A/K48A/K50A (bacterial expression). Used to make protein:<br>2A<sub>-6H</sub> R46A/K48A/K50A | This work | N/A |
| 2A_pOPTnH K73A (bacterial expression). Used to make protein:<br>2A<sub>-6H</sub> K73A | This work | N/A |
| 2A_pOPTnH 9-136, C111S (bacterial expression). Used to make protein: 2A<sub>6H</sub> 9-136; C111S | This work | N/A |
| 2A_pGEX6P1 (cloning intermediate). Used to make SII-2A_pGEX6P1 construct via insertion of annealed oligos at BamHI site (SII_F and SII_R, **Table 5**) | Napthine et al., 2017 | N/A |
| SII-2A_pGEX6P1 (bacterial expression; ORF encoding GST-3C-SII-2A). Used to make protein SII-2A, which is released after removal of GST tag with 3C protease. | This work | N/A |
| pMZ_lambda_OT EMCV CCC | This work | N/A |
| pMZ_lambda_OT EMCV CUC | This work | N/A |
| **Software and Algorithms** |  |  |
| XIA2 | Winter, 2009 | N/A |
| XDS | Kabsch, 2010 | N/A |
| AIMLESS | Evans and Murshudov, 2013 | N/A |
| autoSHARP | Vonrhein et al., 2007 | N/A |
| SHELXD | Sheldrick, 2008 | N/A |
| SHARP |  | N/A |
| SOLOMON | Abrahams and Leslie, 1996 | N/A |
| ARP/wARP | Perrakis et al., 2001 | N/A |
| Phaser | McCoy et al., 2007 | N/A |
| phenix.autobuild | Terwilliger et al., 2008 | N/A |
| COOT | Emsley et al., 2010 | N/A |
| phenix.refine | Adams et al., 2010 | N/A |
| ISOLDE | Croll et al., 2018 | N/A |
| MolProbity | Chen et al., 2010 | N/A |
| PDBeFOLD | Krissinel and Henrick, 2004 | N/A |
| DALI | Holm and Laakso, 2016 | N/A |
| PDBePISA, European Bioinformatics Institute, EBI | Krissinel and Henrick, 2007 | N/A |
| CATHEDRAL | Redfern et al., 2007 | N/A |
| PDB2PQR | Dolinsky et al., 2004 | N/A |
| PROPKA | Li et al., 2005 | N/A |
| APBS | Baker et al., 2001 | N/A |
| EPU | FEI company | N/A |
| MotionCor 2 | Zheng et al., 2017 | N/A |
| CtfFind4 | Rohou and Grigorieff, 2015 | N/A |
| RELION 3.1 | Zivanov et al., 2018 | N/A |
| PyMOL 1.5.0.5 | Schrödinger LLC | N/A |
| ChimeraX | Pettersen et al., 2004 | N/A |
| ImageQuant<sup>TM</sup> TL. 1-D gel electrophoresis, dot blots, arrays, user-defined gel analysis | GE Healthcare | N/A |
| ImageStudio Lite | LI-COR | N/A |
| InkScape 0.92.3 | https://inkscape.org/ | N/A |
| Matlab | https://www.mathworks.com/products/matlab.html | N/A |
| Spyder | https://www.spyder-ide.org/ | N/A |
| Graphpad | https://www.graphpad.com/ | N/A |
| Bluelake | Lumicks | N/A |
| RNAstructure | https://rna.urmc.rochester.edu/RNAstructure.html | N/A |
| SimRNAweb | Magnus et al., 2016 | N/A |
| **Other** |  |  |
| BioRad 4-20 % Mini PROTEAN TGX precast gels | BioRad | 4561096, 4561094 |
| Amicon Ultra Centrifugal Filter Units (10K MMCO, various) | Merck Millipore | UFC901008, UFC801008, UFC501024 |
| S200 HR microspin gel filtration columns | Cytiva | 27-5120-01 |
| Nylon CryoLoops (for crystal mounting) | Hampton | HR4-932, HR4-945, HR4-947 |
| Quantifoil R 2/2 400-mesh copper supports | Electron Microscopy Sciences | Q4100CR2 |
| Streptavidin Coated Polystyrene Particles, 1.5-1.9 μm, 5 ml, 1.0% w/v | Spherotech | SVP-15-5 |
| Anti-digoxigenin Coated Polystyrene Particles, 2.0-2.4 μm, 2 ml, 0.1% w/v | Spherotech | DIGP-20-2 |
| Monolith NT.115 Premium Capillaries | Nanotemper | MO-K025 |
| GE Storage Phosphor Screens | GE Healthcare | BAS-IP MS 2025 |
| Carestream® BioMax® MR film | MERCK | Z350389-50EA |

